# Phosphorylation and pocket-binding ligands rewire the coupled networks in multidomain protein condensate

**DOI:** 10.64898/2026.08.07.743494

**Authors:** Yu Cao, Jinyu Chen, Xiakun Chu

## Abstract

Biomolecular condensates formed by multidomain proteins are increasingly linked to disease and are emerging targets for chemical modulation. Yet most molecular models emphasize intrinsically disordered regions (IDRs), whereas many available ligands bind folded-domain pockets. How IDR modifications and folded-domain ligands are integrated to control condensate organization remains unclear. Here, we address this problem using histone deacetylase 6 (HDAC6), a disease-associated deacetylase whose S22 phosphorylation and catalytic-pocket ligands differentially regulate condensation. Using atomistic-informed Martini 3 coarse-grained simulations, we show that S22 phosphorylation enhances phase separation of the N-terminal IDR1 without compacting isolated chains. Instead, phosphorylation creates a phospho-S22/D26-centered anionic contact node that strengthens dense-phase interactions with Arg/Lys-rich patches. The HDAC6 ligands Nexturastat A and HPOB remodel this IDR network through distinct charged-contact mechanisms: Nexturastat A acts as a focused competitor for cationic patches, whereas HPOB permits partial compensation through alternative charged contacts. In full-length phospho-HDAC6, the assembled network is not IDR1-dominated but is organized by FD3, the C-terminal ZnF-UBP domain. Pocket-bound ligands redirect this folded-domain network from FD3–FD3 association toward pocket–FD3 contacts, with stronger and more persistent remodeling by Nexturastat A. These results reveal a two-layer regulatory mechanism in which phosphorylation rewires IDR electrostatics while pocket-binding ligands redirect folded-domain interactions, suggesting a strategy for modulating multidomain condensates through ligand-induced contact-network remodeling.

## Introduction

Biomolecular condensates concentrate proteins and nucleic acids into membrane-free biochemical compartments, thereby organizing reactions, storage, stress responses and gene regulation through multivalent molecular interactions (Hyman et al., 2014; Shin and Brangwynne, 2017; Banani et al., 2017; Alberti et al., 2019; Boeynaems et al., 2018). Dysregulation of condensate assembly and material properties is increasingly linked to cancer, neurodegeneration and other disease states (Mitrea et al., 2022; Jiang and Kang, 2025; Nam and Gwon, 2023). Many disease-associated condensates are formed by multidomain proteins, including RNA-binding proteins, transcriptional regulators and signaling proteins that combine folded domains with intrinsically disordered regions (IDRs) (Molliex et al., 2015; Guillén-Boixet et al., 2020; Sanders et al., 2020; Martin et al., 2021; Sabari et al., 2018; Hess and Joseph, 2025). How these distinct architectural elements cooperate to build condensates, and how their interactions can be selectively modulated, remains a central question in condensate biology.

Current molecular models of liquid–liquid phase separation have been shaped largely by studies of IDRs, low-complexity sequences, charge-patterned polyampholytes and sticker-and-spacer architectures (Nott et al., 2015; Pak et al., 2016; Martin and Holehouse, 2020; Wang et al., 2018; Martin et al., 2020; Vernon et al., 2018; Dignon et al., 2018; Gilat et al., 2026). This framework has provided powerful principles for understanding how sequence composition and patterning control IDR-driven condensation. Recent systematic mapping of IDR–IDR interactions further suggests that many condensate-forming IDRs can coassemble promiscuously, with phase-separation propensity linked to overall sequence stickiness, whereas discrete condensate identity may require additional structural determinants (Gilat et al., 2026). For multidomain proteins, folded domains are therefore unlikely to act only as passive cargo appended to disordered scaffolds. Instead, they can contribute multivalent binding surfaces, impose architectural constraints, tune the behavior of neighboring disordered regions and create ligand-accessible regulatory nodes (Li et al., 2012; Peran and Mittag, 2020; Hess and Joseph, 2025; Wang and Marrink, 2026; Martin et al., 2021). This raises an important mechanistic and pharmacological possibility: small molecules designed to bind folded-domain pockets may regulate condensates not only by altering enzymatic activity, but also by reshaping domain-level interaction networks.

Histone deacetylase 6 (HDAC6) provides a compelling system in which to test this idea. HDAC6 is a cytoplasmic class IIb deacetylase containing two folded catalytic domains and a C-terminal ubiquitin-binding zinc-finger domain, connected by extended disordered regions as schematized in Figure S1. Beyond histone deacetylation, HDAC6 regulates microtubule acetylation, misfolded- protein handling, aggresome formation, autophagy, stress-granule responses and cancer-associated signaling (Hubbert et al., 2002; Matsuyama et al., 2002; Haggarty et al., 2003; Kawaguchi et al., 2003; Pandey et al., 2007; Kwon et al., 2007; Aldana-Masangkay and Sakamoto, 2011; Pulya et al., 2021). Recent work has connected HDAC6 to phosphorylation-regulated phase separation in triple- negative breast cancer. Phosphorylation of S22 enhances the phase-separation tendency of the N- terminal disordered region IDR1, and S22-phosphorylated full-length HDAC6 forms condensates (Lu et al., 2024). Importantly, deletion of IDR1 reduces but does not abolish condensation of the remaining HDAC6 construct, indicating that regions outside IDR1 also contribute to assembly (Lu et al., 2024). The same study found that Nexturastat A strongly suppresses condensation of both S22-phosphorylated IDR1 and S22-phosphorylated HDAC6, whereas HPOB has a weaker disruptive effect (Lu et al., 2024). HDAC6 therefore links three features that are often considered separately: phosphorylation-dependent IDR interactions, folded-domain organization and ligand-sensitive condensate regulation.

These observations create a mechanistic puzzle. The experimentally identified regulatory modi- fication, S22 phosphorylation, lies within IDR1, whereas Nexturastat A and HPOB were developed as HDAC6-targeting ligands that bind folded catalytic-domain pockets rather than low-complexity disordered regions (Bergman et al., 2012; Lee et al., 2013; Hai and Christianson, 2016; Miyake et al., 2016; Osko et al., 2020). It is therefore unclear whether these ligands perturb HDAC6 condensation by directly competing with IDR1-mediated contacts, by altering folded-domain interactions, or by coupling these two interaction layers. Resolving this question requires a molecular description that can connect phosphorylation-dependent IDR behavior with ligand-dependent reorganization of the full-length multidomain assembly.

Here, we develop an atomistic-informed Martini 3 coarse-grained simulation framework to examine wild-type HDAC6 (WT-HDAC6), S22-phosphorylated HDAC6 (phospho-HDAC6), and the HDAC6-targeting ligands Nexturastat A and HPOB (Souza et al., 2021). The full-length model combines domain organization inferred from per-residue predicted local distance difference test (pLDDT) scores from AlphaFold (Jumper et al., 2021; Varadi et al., 2022), Martini3-IDP representations for disordered regions (Wang et al., 2025), and GōMartini 3 restraints for folded domains (Souza et al., 2025). Because phosphorylation and ligand chemistry are central to the mechanisms studied here, we calibrated a local phosphoserine representation against atomistic reference simulations and generated molecule-specific coarse-grained models of Nexturastat A and HPOB, followed by validation simulations of catalytic-pocket engagement (Linhartova et al., 2024; Szczuka et al., 2026; Graham et al., 2017).

Using this framework, we show that S22 phosphorylation enhances IDR1 phase separation without compacting isolated IDR1 chains. Instead, phosphorylation rewires dense-phase electrostatic contacts by creating phospho-S22-centered interactions with Arg/Lys-rich regions. We further find that Nexturastat A and HPOB perturb IDR1 condensation through distinct charged- contact competition and compensation mechanisms, with Nexturastat A more strongly disrupting the phospho-S22-centered contact network. Finally, full-length simulations reveal that phospho- HDAC6 assembly is not governed by IDR1 alone, but is organized around interactions involving the C-terminal folded domain FD3. Pocket-binding ligands redirect this FD3-centered network from FD3–FD3 association toward pocket–FD3 contacts, with stronger remodeling by Nexturastat A. These results identify folded-domain interaction networks as key organizers of multidomain-protein condensates and suggest that catalytic-pocket ligands can regulate condensate organization by rewiring domain-level contacts.

## Results

An atomistic-informed coarse-grained framework captures HDAC6 modularity and ligand pocket recognition

To investigate how S22 phosphorylation and HDAC6-targeting ligands regulate condensate formation, we first constructed coarse-grained models of full-length HDAC6, Nexturastat A and HPOB. The full-length HDAC6 model preserves the modular organization of the protein, with folded and disordered blocks assigned using the AlphaFold-predicted per-residue pLDDT profile together with the known HDAC6 domain architecture, as detailed in Computational Methods and Figure S2A. This assignment defined three folded domains, FD1, FD2 and FD3, and three disordered regions, IDR1, IDR-L and IDR2 (Figure 1A). IDR1, IDR-L and IDR2 were represented using the Martini3-IDP framework (Wang et al., 2025), whereas FD1, FD2 and FD3 were stabilized with GōMartini 3 restraints (Souza et al., 2025). This design allowed the disordered regions to remain flexible while maintaining the structural integrity of the folded domains.

**Figure 1.**
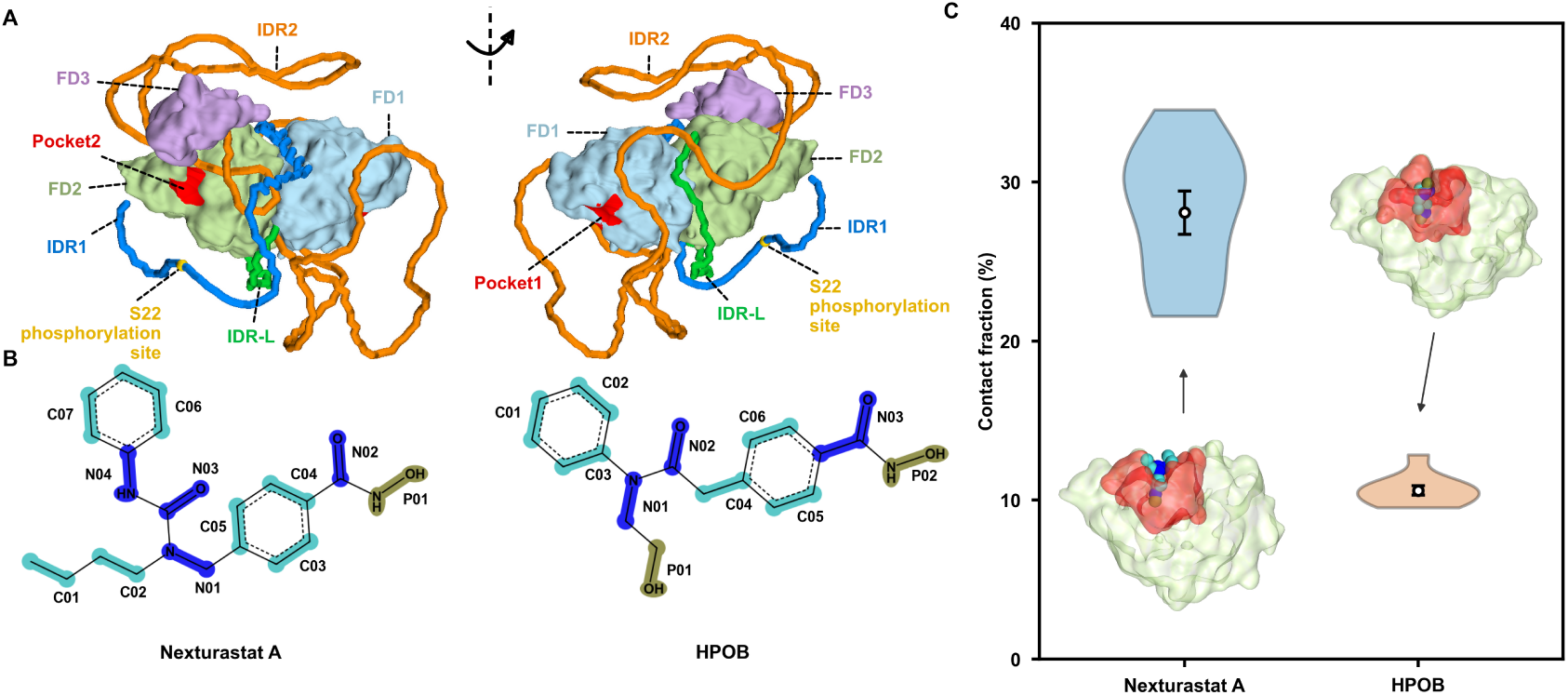
Coarse-grained model construction and ligand pocket-targeting validation for HDAC6. **(A)** Front and rear views of the full-length HDAC6 coarse-grained model. Folded domains are shown as FD1 (light blue), FD2 (light green) and FD3 (light purple), and disordered regions are shown as IDR1 (blue), IDR-L (green) and IDR2 (orange). The S22 phosphorylation site is highlighted in yellow, and pocket1 and pocket2 on the folded catalytic domains are highlighted in red. **(B)** Chemical structures and final Martini 3 coarse-grained mappings of Nexturastat A and HPOB. **(C)** Ligand–pocket contact fractions from reduced FD2–ligand validation simulations. Representative FD2-bound ligand configurations are shown with the FD2 pocket highlighted in red. Violin plots show replicate distributions, and overlaid markers with error bars indicate mean ± s.e.m. across ten independent 10 *µ*s simulations for each ligand. These simulations assess qualitative pocket-recognition behavior and are not binding-affinity calculations.

Because S22 phosphorylation is the experimentally identified modification that enhances HDAC6 IDR1 condensation, we next refined its local coarse-grained representation. Following a previously reported Martini 3 approximation for phosphoserine (Linhartova et al., 2024), the S22 side-chain bead was assigned as Q5 with a charge of −1*e*, while the remaining protein bead assignment was kept unchanged. To improve the local geometry around the phosphorylated residue, bonded parameters involving the S22 side-chain bead, including bond, angle, dihedral and improper terms, were calibrated against mapped all-atom reference trajectories (Figure S2B–H). In the full-length model, the S22 phosphorylation site is located within IDR1, whereas pocket1 and pocket2 are defined as ligand-accessible regions on the two folded catalytic domains (Figure 1A).

We then generated molecule-specific Martini 3 models of Nexturastat A and HPOB (Figure 1B). Both compounds have been reported as potent HDAC6 inhibitors (Bergman et al., 2012; Lee et al., 2013), and inhibitor-bound HDAC6 catalytic-domain structures have established the structural basis of catalytic-pocket recognition and HDAC6 selectivity (Hai and Christianson, 2016; Miyake et al., 2016; Osko et al., 2020). Each ligand was represented as a neutral Martini 3 small molecule. Initial atom-to-bead mappings and bead types were generated from the all-atom ligand structures, and bonded geometries were refined against mapped atomistic ligand trajectories (Szczuka et al., 2026; Graham et al., 2017). This procedure retained compound-specific polar and hydrophobic bead patterns, allowing subsequent simulations to compare ligand-specific interaction modes rather than generic small-molecule effects.

We next tested whether the resulting ligand models retained qualitative recognition of the catalytic pocket after coarse graining. For this purpose, we performed reduced validation simulations containing one FD2 domain and one ligand molecule. Because FD1 and FD2 share the catalytic-domain architecture of HDAC6, pocket2 on FD2 was used as a representative catalytic pocket for these controls. Ten independent 10 *µ*s simulations were performed for each ligand, and ligand– pocket engagement was quantified as the fraction of trajectory frames in which the ligand contacted the pocket. Both ligands showed intermittent pocket engagement across replicate trajectories, but the contact intervals were denser for Nexturastat A than for HPOB (Figure S3). Consistently, the mean pocket-contact fraction was higher for Nexturastat A than for HPOB, 28.1 ± 1.4% versus 10.6 ± 0.3%, respectively, across ten independent simulations (Figure 1C). Representative bound configurations illustrate that both coarse-grained ligands can access the intended FD2 pocket.

Together, the full-length HDAC6 model, the calibrated phospho-S22 representation, the compound-specific ligand models and the FD2–ligand pocket-recognition controls provide the structural basis for analyzing how phosphorylation and ligand binding reshape HDAC6 condensate-forming interactions. We therefore next examined how S22 phosphorylation alters the phase separation of the isolated IDR1 segment.

### S22 phosphorylation promotes IDR1 phase separation by rewiring dense-phase electrostatic contacts

Previous experiments showed that S22 phosphorylation enhances the condensation tendency of the isolated N-terminal disordered region IDR1 relative to wild-type IDR1 (WT-IDR1) (Lu et al., 2024). To connect our simulations directly to this experimental construct, the isolated-IDR1 simulations in Figure 2 used residues 1–66, matching the experimentally defined IDR1 fragment. This construct is shorter than the pLDDT-defined IDR1 block used in the full-length HDAC6 model, which spans residues 1–85. The full-length HDAC6 sequence is provided in Figure S4.

**Figure 2.**
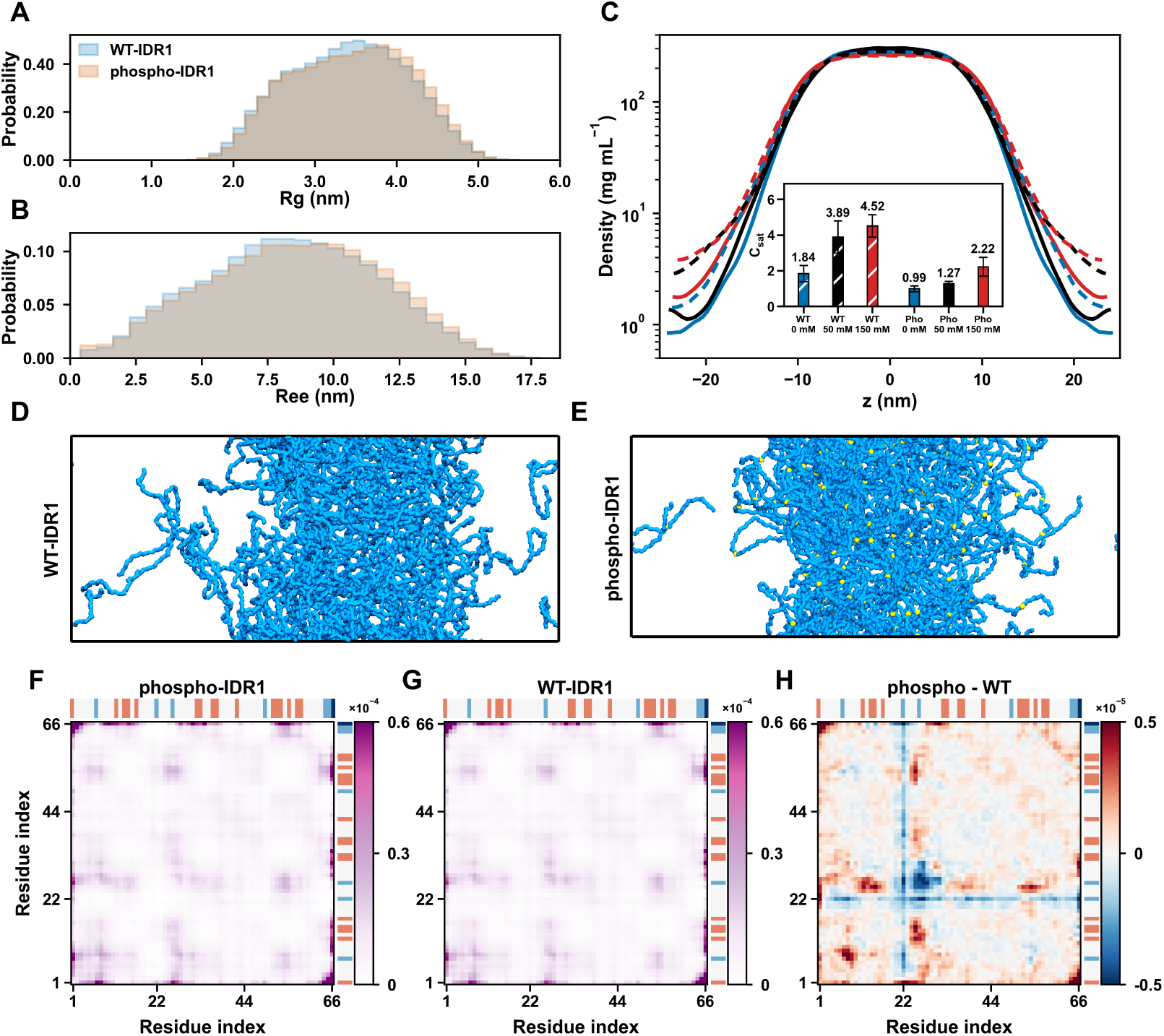
S22 phosphorylation enhances IDR1 phase separation without promoting single-chain compaction and rewires dense-phase interchain contacts. **(A,B)** Radius-of-gyration (*R_g_*) and end-to-end distance (*R*_ee_) distributions from single-chain simulations of residues 1–66 WT-IDR1 and phospho-IDR1 at 50 mM NaCl. **(C)** Protein density profiles along the slab-normal *z* axis from WT-IDR1 and phospho-IDR1 direct-coexistence simulations at 0, 50 and 150 mM NaCl. Profiles were calculated over the 11–15 *µ*s analysis window and are shown on a logarithmic scale. Blue, black and red curves indicate 0, 50 and 150 mM NaCl, respectively; dashed and solid curves indicate WT-IDR1 and phospho-IDR1, respectively. The inset shows the apparent dilute-phase concentration, *C*_sat_, estimated from the dilute regions of the same density profiles. Error bars indicate dilute-region variability from the density-analysis procedure. **(D,E)** Representative 15 *µ*s snapshots from WT-IDR1 and phospho-IDR1 slab simulations, cropped consistently and displayed at matched scale. **(F,G)** Dense-phase interchain residue–residue contact probability maps for phospho-IDR1 and WT-IDR1 at 50 mM NaCl, calculated over the 11–15 *µ*s analysis window and shown using the same color scale. **(H)** Difference contact map calculated as phospho-IDR1 minus WT-IDR1. Red indicates contacts enhanced by S22 phosphorylation, whereas blue indicates contacts reduced by phosphorylation. Charge strips above and to the right of each map indicate residue-level charge states using a discrete encoding: red, +1; white, 0; light blue, −1; dark blue, −2.

We first asked whether phosphorylation-enhanced condensation could be explained by compaction of individual IDR1 chains. Single-chain simulations at 50 mM NaCl showed strongly overlapping radius-of-gyration distributions for WT-IDR1 and phospho-IDR1 (Figure 2A). Rather than shifting toward a more compact ensemble, phospho-IDR1 sampled slightly larger *R_g_* values. The end-to- end distance distribution showed the same tendency, with phospho-IDR1 sampling marginally more extended conformations than WT-IDR1 (Figure 2B). Consistently, replicate-level mean values showed a small increase in end-to-end distance from 8.33 ± 0.18 nm for WT-IDR1 to 8.56 ± 0.21 nm for phospho-IDR1, and a small increase in *R_g_* from 3.40 ± 0.04 nm to 3.45 ± 0.04 nm (Figure S5A,B). Pair-distance distribution functions were also nearly superimposable and showed no shift toward shorter intrachain distances upon phosphorylation (Figure S5C). Thus, S22 phosphorylation does not enhance IDR1 phase separation by compacting the dilute single-chain ensemble.

We therefore examined whether phosphorylation instead alters intermolecular interactions in the condensed phase. Direct-coexistence slab simulations of WT-IDR1 and phospho-IDR1 were performed at 0, 50 and 150 mM NaCl. Both constructs formed coexisting dense and dilute phases under all salt conditions examined (Figure 2C–E). However, phospho-IDR1 consistently showed a lower apparent dilute-phase concentration, *C*_sat_, than WT-IDR1. For WT-IDR1, the estimated *C*_sat_ values were 1.84, 3.89 and 4.52 mg ml^−1^ at 0, 50 and 150 mM NaCl, respectively. The corresponding values for phospho-IDR1 were 0.99, 1.27 and 2.22 mg ml^−1^ (Figure 2C). Because a lower *C*_sat_ indicates stronger phase separation, these slab simulations recapitulate the experimentally observed enhancement of IDR1 condensation by S22 phosphorylation (Lu et al., 2024). Increasing NaCl raised *C*_sat_ for both constructs, indicating that electrostatic interactions contribute substantially to IDR1 phase separation. Consistent with this interpretation, the salt-dependent contact-map analysis showed that phosphorylation-enhanced contacts involving the S22-containing region and positively charged IDR1 clusters were strongest at low salt and attenuated at higher salt (Figure S6).

To identify the intermolecular contacts underlying the stronger phase separation of phospho- IDR1, we calculated dense-phase interchain residue–residue contact maps from the 50 mM NaCl slab simulations. High-probability contacts were not distributed uniformly along the chain. Instead, they were concentrated around charge-enriched sequence regions, including the N-terminal Arg-rich region around residues 12–18, the S22/D26-containing region around residues 21–26, the central Lys/Arg-rich region around residues 32–42 and the C-terminal Lys-rich region around residues 51–60 (Figure 2F,G). Thus, IDR1 condensation is organized by sequence-patterned electrostatic contacts among charged patches rather than by nonspecific collapse of the entire chain.

The phospho-minus-WT difference map revealed how S22 phosphorylation rewires this densephase contact network (Figure 2H). The most prominent phosphorylation-enhanced features were off-diagonal contacts connecting the S22/D26-containing region with positively charged segments on other chains. These included contacts with the N-terminal Arg-rich cluster around residues 12–18 and the C-terminal Lys-rich cluster around residues 51–60. This pattern indicates that the added phosphate at S22 creates a localized anionic interaction site that can engage spatially separated cationic patches in trans. In contrast, several WT-like contacts involving the S22-proximal region were reduced after phosphorylation, particularly contacts that lacked the same charge-complementary character. Phosphorylation therefore does not simply increase the total number of IDR1 contacts. Instead, it redistributes the dense-phase interaction network, selectively strengthening phospho- S22/D26-to-Arg/Lys contacts while weakening alternative contacts involving the same local region. Together, these results reveal an apparent decoupling between dilute-chain compaction and phase-separation propensity. S22 phosphorylation slightly expands, rather than compacts, isolated IDR1 chains, yet it strengthens phase separation by creating new charge-complementary interchain contacts in the dense phase. The phosphorylation-enhanced condensate is therefore stabilized by electrostatic rewiring centered on the phospho-S22/D26-containing segment and Arg/Lys-rich IDR1 patches. This mechanism provides a molecular explanation for the experimentally observed enhancement of IDR1 condensation upon S22 phosphorylation and establishes the charged-contact network that we next use to interpret ligand-dependent perturbation of IDR1 condensates.

### Nexturastat A and HPOB modulate IDR1 condensation by redistributing charged contacts

The phosphorylation analysis above showed that IDR1 condensation is stabilized by sequence- patterned electrostatic contacts between acidic segments and Arg/Lys-rich clusters, with phospho- S22 creating an additional anionic contact node. We therefore next asked whether HDAC6-targeting ligands perturb IDR1 condensation by redistributing this charged interchain contact network. To test this, we performed 50 mM NaCl direct-coexistence slab simulations of WT-IDR1 and phospho-IDR1 in the presence of either Nexturastat A or HPOB and analyzed the 11–15 *µ*s production window.

Slab-density profiles revealed a ligand response that depended strongly on phosphorylation state (Figure 3A). In WT-IDR1, the apparent dilute-phase concentration, *C*_sat_, was 3.89 mg ml^−1^ in the ligand-free system. Nexturastat A increased this value to 7.81 mg ml^−1^, consistent with weakened phase separation, whereas HPOB decreased it to 0.92 mg ml^−1^, suggesting stronger apparent condensation. In phospho-IDR1, the ligand-free *C*_sat_ was 1.27 mg ml^−1^. Nexturastat A produced a larger increase to 4.58 mg ml^−1^, and HPOB increased this value to 2.17 mg ml^−1^. Thus, both ligands weakened phospho-IDR1 phase separation, but Nexturastat A produced the stronger effect, increasing *C*_sat_ by approximately 3.6-fold relative to ligand-free phospho-IDR1, compared with approximately 1.7-fold for HPOB. Representative phospho-IDR1 slab snapshots are shown in Figure 3B, and the corresponding WT-IDR1 snapshots are provided in Figure S7.

**Figure 3.**
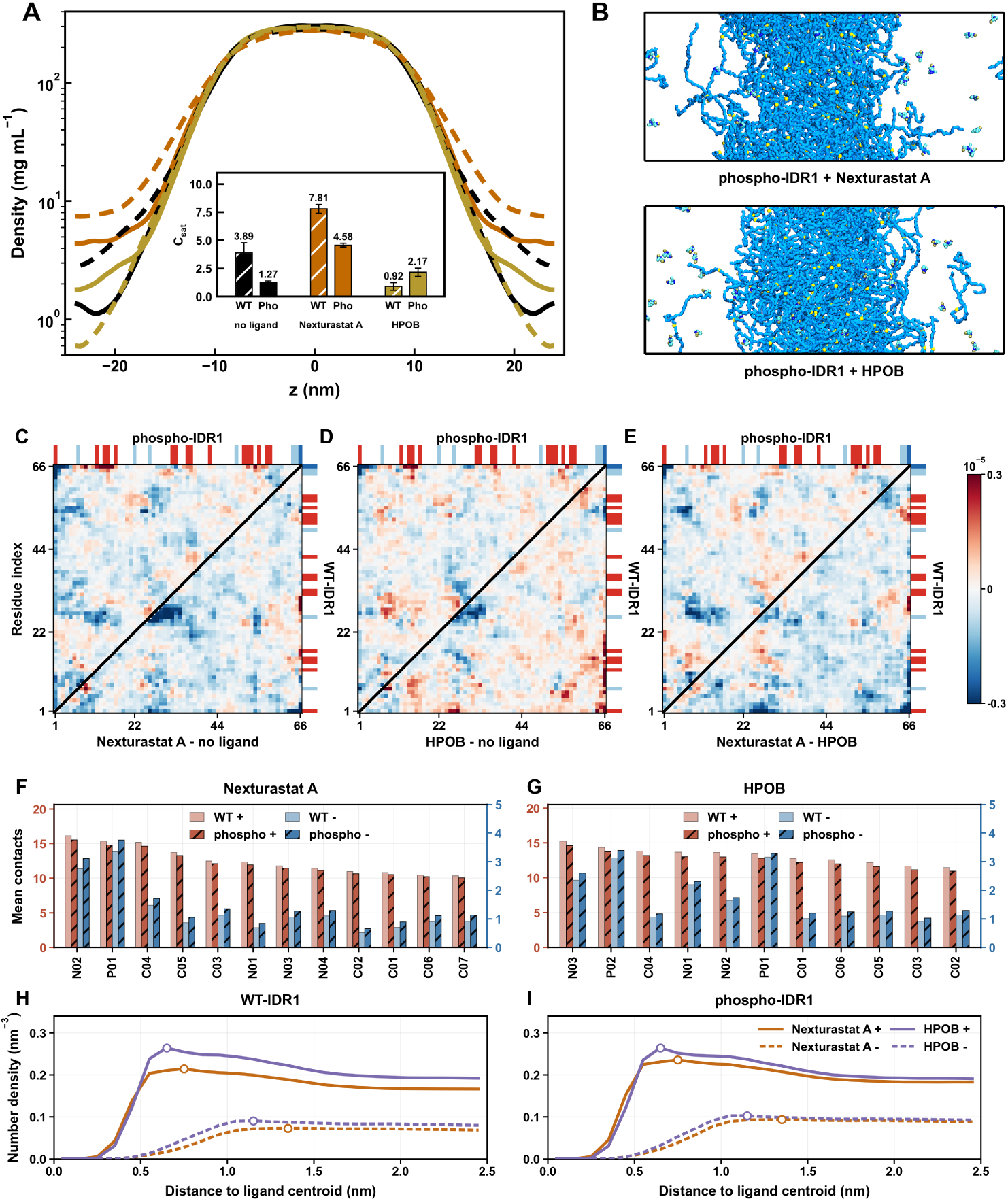
Nexturastat A and HPOB differentially perturb IDR1 condensation through charged-contact reorganization. **(A)** Slab density profiles for WT-IDR1 and phospho-IDR1 at 50 mM NaCl under ligand-free, Nexturastat A-treated and HPOB-treated conditions, calculated over the 11–15 *µ*s analysis window. Dashed and solid lines indicate WT-IDR1 and phospho-IDR1, respectively. Curve colors match the inset bar groups. The inset shows the apparent dilute-phase concentration, *C*_sat_, estimated from the same density profiles. **(B)** Representative 15 *µ*s snapshots from phospho-IDR1 slabs treated with Nexturastat A and HPOB, shown from top to bottom. **(C–E)** Triangular dense-phase interchain residue–residue contact difference maps. Upper and lower triangles show phospho-IDR1 and WT-IDR1, respectively. The comparisons are Nexturastat A minus no ligand, HPOB minus no ligand, and Nexturastat A minus HPOB. Red and blue indicate contacts increased and reduced in the first condition of each comparison, respectively. Charge strips indicate residue-level charge states: red, +1; white, 0; light blue, −1; dark blue, −2. **(F,G)** Ligand-bead contacts with positively and negatively charged IDR1 residues for Nexturastat A and HPOB. In the panel labels, + and − denote positively and negatively charged residues, respectively. **(H,I)** Radial number-density profiles of positively and negatively charged IDR1 residues around ligand centroids in WT-IDR1 and phospho-IDR1 dense phases. Orange and purple curves indicate Nexturastat A and HPOB, respectively; solid and dashed lines indicate positively and negatively charged residues, respectively. Circle markers indicate profile peak positions.

We next examined whether these density changes were associated with ligand-dependent changes in dense-phase interchain contacts. The triangular contact-difference maps in Figure 3C–E compare ligand-treated and ligand-free systems, with phospho-IDR1 shown in the upper triangles and WT-IDR1 in the lower triangles. For interpretation, we focused on the major charged regions in the residues 1–66 IDR1 construct: D7, the N-terminal Arg-rich patch around residues 12–18, the S22/D26-containing region around residues 21–26, the central Lys/Arg-rich patch around residues 32–42, E49, the C-terminal Lys-rich patch around residues 51–58, and the acidic C-terminal tail around residues 64–66. The corresponding sequence context is provided in Figure S4.

Nexturastat A caused a broad but structured redistribution of dense-phase contacts in phospho- IDR1 (Figure 3C). The most coherent contact losses were centered on the S22/D26-containing region and on its off-diagonal contacts with the N-terminal Arg-rich patch, the central Lys/Arg-rich patch, and, more weakly, the C-terminal Lys-rich patch. These are the same acidic-to-basic contacts that were enhanced by phosphorylation in Figure 2H, indicating that Nexturastat A preferentially weakens the phospho-S22/D26-centered electrostatic network that stabilizes phospho-IDR1 condensation. Additional contact changes extended into the N-terminal, central and C-terminal charged regions, showing that Nexturastat A does not eliminate a single residue-pair interaction. Rather, it remodels the dense-phase contact landscape, with the clearest effect being depletion of phospho-S22/D26-to-Arg/Lys-rich contacts. In WT-IDR1, Nexturastat A also reduced contacts involving the D26-containing region and nearby basic patches, but these changes were broader and less organized than in phospho-IDR1, consistent with the absence of the added phospho-S22 anionic site.

HPOB produced a more mixed response (Figure 3D). In phospho-IDR1, HPOB reduced part of the S22/D26-centered network, most notably contacts between the S22/D26-containing region and the central Lys/Arg-rich patch. However, these losses were less coherent and less extensive than those induced by Nexturastat A. HPOB also increased contacts involving other charged regions, including the N-terminal charged region, the S22-proximal region, the C-terminal Lys-rich patch and the acidic C-terminal tail. Thus, HPOB partially competes with the phospho-S22-centered electrostatic network but also preserves or replaces some contacts through alternative charged-region interactions. This mixed response explains why HPOB weakened phospho-IDR1 phase separation more modestly than Nexturastat A.

The WT-IDR1 response further illustrates this compensation. In the lower triangle of Figure 3D, HPOB generated multiple positive contact changes involving the N-terminal charged region, the D26-containing region, the central basic patch and C-terminal charged/basic segments. Because WT-IDR1 lacks the strong phospho-S22-centered interaction node, these HPOB-associated alternative contacts can compensate for, and in this case outweigh, the loss of native contacts. This provides a contact-level explanation for the decrease in WT-IDR1 *C*_sat_ from 3.89 to 0.92 mg ml^−1^ in the presence of HPOB.

The direct Nexturastat A-minus-HPOB comparison highlights the ligand-specific nature of this contact redistribution (Figure 3E). In phospho-IDR1, contacts were lower with Nexturastat A than with HPOB around the S22/D26-containing region and its interactions with the N-terminal Arg-rich and central Lys/Arg-rich patches, with additional losses extending into C-terminal charged/basic contacts. In contrast, contact gains with Nexturastat A were more scattered and did not form a coherent phospho-S22/D26-centered acidic-to-basic pattern. Thus, the main difference between the two ligands is not that Nexturastat A globally removes all residue–residue contacts. Instead, Nexturastat A more strongly depletes the phosphorylation-enhanced acidic-to-basic contact network, whereas HPOB retains more charged-region contacts. In WT-IDR1, HPOB also retained or enhanced more contacts involving the D26-containing segment and C-terminal charged/basic patches, consistent with its stronger apparent condensation effect in the WT system.

To understand the molecular origin of these different contact-map responses, we analyzed ligand-bead contacts with charged IDR1 residues. For Nexturastat A, contacts with positively charged residues were concentrated at the N02–P01 polar end of the molecule (Figures 1B, 3F and Figure S8). Contacts with negatively charged residues were much weaker, indicating preferential association with Arg/Lys-rich patches. This focused contact mode provides a mechanistic explanation for the contact-map changes: by occupying cationic IDR1 patches, Nexturastat A reduces their availability for interchain contacts with phospho-S22, D26 and other acidic residues.

HPOB showed a related but less localized contact pattern. Its N03–P02 hydroxamate/amide-associated end contacted positively charged residues, but HPOB also retained an additional lateral P01 polar site that contributed to charged-residue contacts (Figures 1B, 3G and Figure S8). These bead-level contacts should be interpreted as coarse-grained polar or electrostatic contacts rather than atomistically resolved directional hydrogen bonds, as summarized in Table S1. The more distributed polar-contact pattern of HPOB provides a plausible basis for its compensatory behavior: HPOB can compete with native anion–cation protein contacts, but its additional polar site allows alternative local contacts with charged IDR1 regions. This compensation strengthens the weaker WT- IDR1 condensate but is insufficient to preserve the more specific phospho-S22-reinforced contact network in phospho-IDR1.

Ligand-centered radial number-density profiles further support this competition-and- compensation model (Figure 3H,I). In both WT-IDR1 and phospho-IDR1 dense phases, positively charged residues were enriched closest to the ligand centroid, with local maxima at approximately 0.6–0.8 nm. Negatively charged residues had lower densities and peaked farther from the ligands, indicating that both compounds preferentially reside near Arg/Lys-rich patches rather than near anionic residues. In phospho-IDR1, however, the negative-residue distributions differed between the two ligands. Around HPOB, negatively charged residues formed a more defined outer shell near 1.1 nm, whereas the corresponding Nexturastat A profile was lower and broader, with a maximum extending around 1.1–1.3 nm. This more pronounced anionic shell around HPOB is consistent with its additional polar-contact mode and with the compensatory charged contacts observed in the contact maps. By contrast, the Nexturastat A profiles are consistent with stronger local enrichment of cationic residues without an equivalent compensatory anionic shell.

Together, these results show that ligand effects on IDR1 condensation depend on both phosphorylation state and ligand-specific contact geometry. Nexturastat A acts primarily as a focused competitor for Arg/Lys-rich contact sites, thereby depleting the acidic-to-basic interactions that stabilize both WT-IDR1 and phospho-IDR1 condensates. HPOB also competes for cationic patches, but its broader polar-contact pattern allows partial compensation through alternative charged contacts. This compensation strengthens the weaker WT-IDR1 condensate but only partially offsets disruption of the phospho-S22-centered network in phospho-IDR1. Thus, ligand regulation of IDR1 condensation is determined not simply by ligand partitioning into the dense phase, but by how each ligand redistributes the specific charged contacts that stabilize the condensate.

If this phospho-IDR1 mechanism were sufficient to explain the full-length HDAC6 phenotype, full-length phospho-HDAC6 assemblies would be expected to remain primarily IDR1-driven. We therefore next tested this prediction by analyzing the domain-level interaction network of full-length HDAC6 assemblies.

### Pocket-binding ligands redirect an FD3-centered contact network in full-length phospho-HDAC6

The IDR1 slab simulations revealed a sequence-level mechanism in which S22 phosphorylation strengthens charge-complementary IDR1 contacts, whereas Nexturastat A and HPOB perturb these contacts by engaging charged IDR1 regions. Full-length HDAC6, however, contains multiple folded domains connected by extended disordered regions, and its assembly organization may therefore involve interaction layers beyond IDR1. To test this possibility, we simulated 15-chain full-length HDAC6 systems under WT and S22-phosphorylated conditions, with or without Nexturastat A or HPOB. Because S22-phosphorylated HDAC6 is the experimentally relevant phase-separating state, we focus on phospho-HDAC6 in the main text and use WT-HDAC6 simulations as reference controls. These finite-size full-length simulations were designed to resolve contact-network organization within associated multichain assemblies rather than to determine full phase boundaries.

We first monitored total interchain contacts during the 10 *µ*s simulations. Starting from dispersed configurations, all systems formed contact-rich multichain assemblies (Figure 4A). In representative trajectories, interchain contacts rose rapidly during the first 1–2 *µ*s, reached approximately 2,000–3,000 contacts, and then increased more gradually before fluctuating within a broad late-time range of approximately 4,000–5,000 contacts. Ligand-free phospho-HDAC6 transiently reached higher contact numbers, approaching 5,000–5,500 contacts, whereas Nexturastat A-treated phospho-HDAC6, HPOB-treated phospho-HDAC6 and ligand-free WT-HDAC6 showed largely overlapping late-time contact ranges. Additional replicates showed the same qualitative tendency to form associated multichain assemblies, although early growth rates and late-time contact numbers varied among simulations (Figure S9). Endpoint snapshots also showed broadly similar compact multichain morphologies across WT and phospho-HDAC6 systems, with and without ligands (Figure S10). Thus, total contact number and endpoint morphology alone were insufficient to explain ligand-dependent regulation. We therefore analyzed the final 3 *µ*s of the trajectories to resolve the internal domain-level organization of the assembled states.

**Figure 4.**
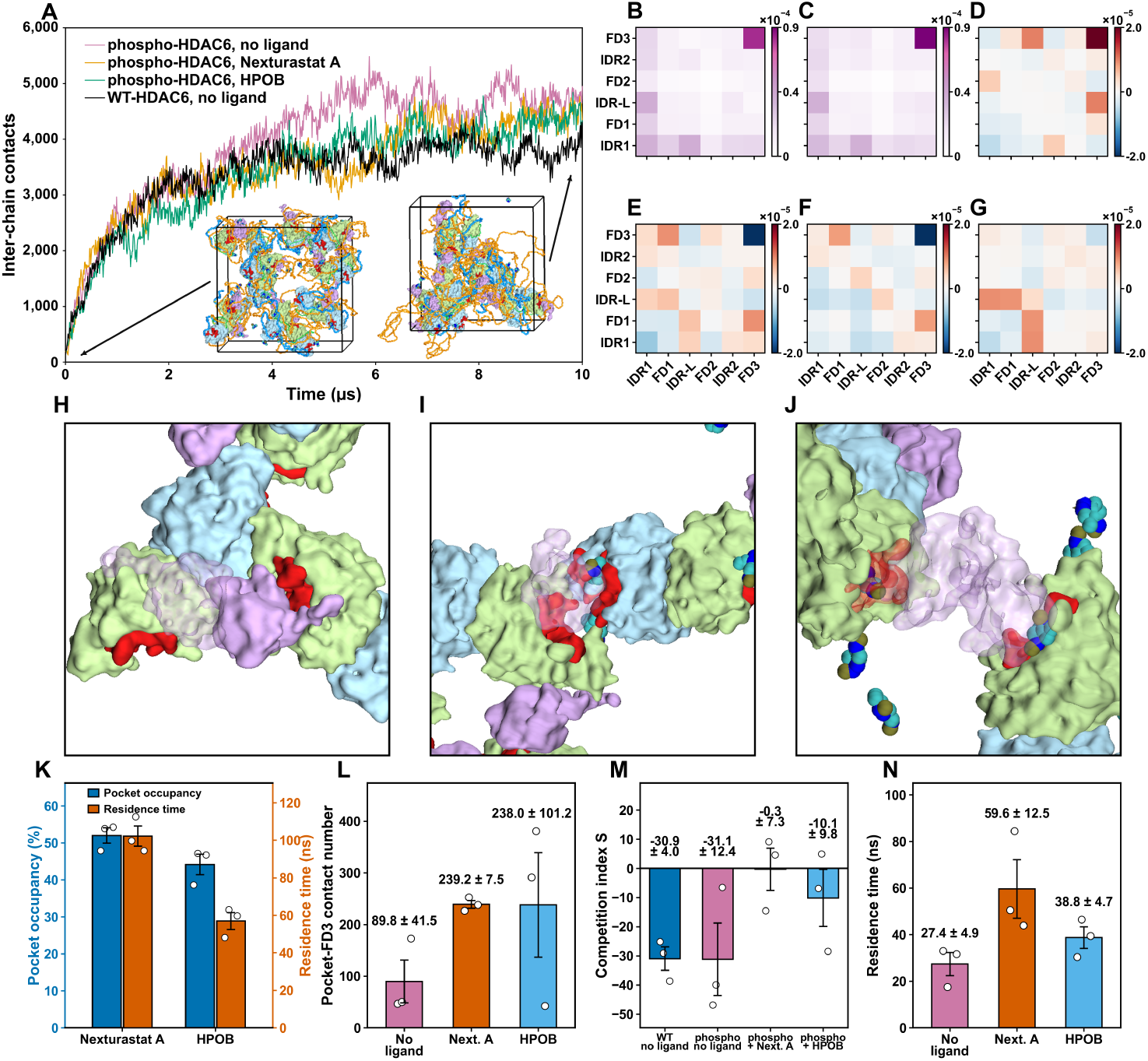
Full-length phospho-HDAC6 assemblies show FD3-centered contact organization and ligand-dependent pocket–FD3 redirection. **(A)** Representative interchain contact-number traces for ligand-free phospho-HDAC6, phospho-HDAC6 with Nexturastat A, phospho-HDAC6 with HPOB, and ligand-free WT-HDAC6 over 10 *µ*s simulations. Embedded snapshots show representative phospho-HDAC6 assembly states at the indicated time points. **(B,C)** Domain-level interchain contact probability maps for ligand-free WT-HDAC6 and ligand-free phospho-HDAC6, respectively, calculated from the final 3 *µ*s analysis window. **(D)** Difference map calculated as ligand-free phospho-HDAC6 minus ligand-free WT-HDAC6. **(E,F)** Difference maps calculated as phospho-HDAC6 with Nexturastat A minus ligand-free phospho-HDAC6 and phospho-HDAC6 with HPOB minus ligand-free phospho-HDAC6, respectively. **(G)** Difference map calculated as phospho-HDAC6 with Nexturastat A minus phospho-HDAC6 with HPOB. In difference maps, red and blue indicate contacts increased and reduced in the first condition of each comparison, respectively. **(H)** Representative local snapshot showing FD3–FD3 contact in ligand-free phospho-HDAC6. **(I,J)** Representative local snapshots showing pocket–FD3 contact in phospho-HDAC6 treated with Nexturastat A and HPOB, respectively. In J, part of FD3 is shown as a semi-transparent surface to expose the contact geometry. **(K)** Mean pocket occupancy and pocket residence time for Nexturastat A and HPOB in full-length phospho-HDAC6 simulations. **(L)** Mean contact number between the combined catalytic-pocket regions, pocket1 and pocket2, and FD3 in ligand-free and ligand-treated phospho-HDAC6 simulations. **(M)** Competition index *S*, calculated from pocket–FD3 and FD3–FD3 contact numbers. More positive values indicate increased pocket–FD3 contacts relative to FD3–FD3 contacts. **(N)** Pocket–FD3 residence time in ligand-free and ligand-treated phospho-HDAC6 simulations. Bars show mean ± s.e.m. across three independent simulations, and open circles indicate individual simulations.

Domain-level interchain contact maps were calculated using the pLDDT-defined blocks IDR1, FD1, IDR-L, FD2, IDR2 and FD3. In both ligand-free WT-HDAC6 and ligand-free phospho-HDAC6, the highest contact frequency occurred in the FD3–FD3 block (Figure 4B,C). Thus, the C-terminal folded domain FD3, corresponding to the ZnF-UBP domain, is a major organizing element of the fulllength assembly. The phospho-minus-WT difference map further showed that S22 phosphorylation increased FD3–FD3 contact frequency and altered additional contacts involving FD3 and other domain blocks, whereas several IDR1-associated contacts were reduced (Figure 4D). Therefore, S22 phosphorylation does not simply amplify an IDR1-only interaction network in the full-length protein. Instead, phosphorylation shifts the multidomain assembly toward stronger FD3-centered organization.

We next asked how ligand treatment alters this FD3-centered contact network. Relative to ligand-free phospho-HDAC6, both Nexturastat A and HPOB reduced FD3–FD3 contact frequency (Figure 4E,F). This was the clearest domain-level contact loss in both ligand-treated systems. At the same time, both ligands increased contacts between FD3 and other regions, including the pocket-containing catalytic domains FD1 and FD2. Because pocket1 and pocket2 reside in FD1 and FD2, respectively, these contact gains indicate that ligand-bound pocket regions provide alternative interaction sites for FD3. Representative local snapshots support this interpretation: ligand-free phospho-HDAC6 showed direct FD3–FD3 association, whereas ligand-treated phospho-HDAC6 showed FD3 positioned near catalytic-pocket regions (Figure 4H–J). Thus, ligand treatment does not globally suppress all domain contacts. Rather, it redirects the full-length contact network away from the original FD3–FD3 mode and toward pocket-associated FD3 contacts.

The direct Nexturastat A-minus-HPOB comparison revealed the ligand-specific component of this redirection (Figure 4G). Nexturastat A produced lower FD3–FD3 contact frequency than HPOB, indicating stronger disruption of the original FD3–FD3 association mode. In contrast, Nexturastat A showed higher contact frequencies in several alternative domain-pair regions, particularly those involving IDR1, IDR-L and FD1. This pattern indicates that Nexturastat A does not simply reduce the total number of interchain contacts. Instead, it more effectively redistributes contacts away from FD3–FD3 association and toward alternative interactions within the multidomain assembly.

We then quantified ligand engagement at the folded-domain pockets in full-length phospho- HDAC6. Pocket occupancy was averaged over pocket1 and pocket2 across all chains and analyzed frames. Nexturastat A showed higher mean pocket occupancy than HPOB, 52.0 ± 2.1% versus 44.2 ± 2.8%, respectively (Figure 4K). The difference was more pronounced for pocket residence time: Nexturastat A remained associated with the pocket regions for 102.2 ± 5.4 ns on average, whereas HPOB showed a shorter mean residence time of 56.9 ± 4.5 ns. Thus, both ligands engage the folded catalytic pockets in full-length assemblies, but Nexturastat A does so more persistently.

Consistent with the contact-map changes, ligand treatment increased direct contacts between catalytic-pocket regions and FD3. The mean (pocket1 + pocket2)–FD3 contact number increased from 89.8±41.5 in ligand-free phospho-HDAC6 to 239.2±7.5 with Nexturastat A and 238.0±101.2 with HPOB (Figure 4L). Thus, both ligands recruited FD3 toward pocket-containing regions to a similar average extent. We next asked whether this increase occurred at the expense of the FD3–FD3 contacts that dominate the ligand-free assembly. To quantify this balance, we calculated the competition index *S*, defined in Computational Methods as a log-scaled ratio comparing pocket–FD3 contacts with FD3–FD3 contacts. More positive values indicate that pocket–FD3 contacts become more prominent relative to FD3–FD3 contacts. In ligand-free WT-HDAC6 and ligand-free phospho-HDAC6, *S* was strongly negative, with values of −30.9 ± 4.0 and −31.1 ± 12.4, respectively (Figure 4M). Ligand treatment shifted *S* to less negative values, reaching −0.3 ± 7.3 with Nexturastat A and −10.1 ± 9.8 with HPOB. This shift indicates that ligand-induced pocket–FD3 contacts compete with FD3–FD3 organization, with the strongest redirection observed for Nexturastat A.

Residence-time analysis further distinguished the two ligands. The mean pocket–FD3 residence time increased from 27.4 ± 4.9 ns in ligand-free phospho-HDAC6 to 59.6 ± 12.5 ns with Nexturastat A and 38.8 ± 4.7 ns with HPOB (Figure 4N). Thus, although Nexturastat A and HPOB generated comparable mean pocket–FD3 contact numbers, Nexturastat A supported longer-lived pocket–FD3 interactions and shifted the competition index closest to zero. The stronger effect of Nexturastat A therefore arises not from a larger average number of pocket–FD3 contacts, but from more persistent pocket-mediated competition against FD3–FD3 association.

Additional analyses supported this domain-redirection mechanism. In two-chain phospho- HDAC6 simulations, the chains remained associated under ligand-free and ligand-treated conditions, but ligand binding altered the pairwise balance between interchain FD3–FD3 association and pocket-centered FD3 recruitment. Consistent with the 15-chain simulations, Nexturastat A produced a higher pocket–FD3 competition index than HPOB (Figure S11). WT-HDAC6 control simulations showed that ligand pocket engagement and pocket–FD3 contacts can occur outside the phosphorylated state, but the redirection pattern was less coherent than in phospho-HDAC6 (Figure S12). In WT-HDAC6, Nexturastat A showed higher mean pocket occupancy and longer mean pocket residence time than HPOB, 49.3 ± 3.8% and 107.3 ± 12.1 ns versus 37.0 ± 2.2% and 46.6 ± 2.0 ns, respectively (Figure S13). Ligand treatment also increased the mean pocket–FD3 contact number relative to ligand-free WT-HDAC6, from 80.4 ± 39.3 to 145.7 ± 39.6 with Nexturastat A and 129.5 ± 40.7 with HPOB (Figure S14). However, the competition index remained negative in both ligand-treated WT systems, reaching −20.8 ± 7.5 with Nexturastat A and −14.9 ± 7.0 with HPOB, and pocket–FD3 residence times showed only modest and variable changes relative to ligand-free WT-HDAC6. These controls indicate that catalytic-pocket engagement is not sufficient by itself to produce the full redirection phenotype; coherent pocket-mediated remodeling of the FD3-centered network is most pronounced in phospho-HDAC6.

Together, the full-length simulations reveal a domain-level mechanism that complements the IDR1 electrostatic mechanism. In isolated IDR1, S22 phosphorylation enhances condensation by rewiring sequence-specific charged contacts. In full-length phospho-HDAC6, however, the dominant organizing feature is FD3–FD3 association, demonstrating that assembly organization cannot be inferred from IDR1 behavior alone. Pocket-binding ligands redirect this FD3-centered folded-domain network by increasing pocket–FD3 contacts and reducing the original FD3–FD3 contact mode. This redirection is more effective for Nexturastat A, which shows higher pocket occupancy, longer pocket residence time, longer pocket–FD3 residence time and a competition index closest to zero. These results identify FD3-centered folded-domain interactions as key organizers of full-length phospho- HDAC6 assemblies and suggest that catalytic-pocket ligands can regulate multidomain condensate organization by redirecting competing domain-level contact modes.

## Discussion

This study identifies a two-layer mechanism by which phosphorylation and pocket-binding ligands regulate phospho-HDAC6 condensate organization. Rather than acting through a single IDR- centered interaction grammar, phospho-HDAC6 assembly is controlled by coupled residue-level and domain-level contact networks. At the sequence level, S22 phosphorylation rewires electrostatic interactions within IDR1 condensates. At the full-length protein level, the dominant organizing feature is an FD3-centered folded-domain network that can be redirected by ligands bound to the catalytic-domain pockets. These findings place HDAC6 within an emerging view of multidomain- protein condensation in which IDRs provide important phase-separation propensity, but folded domains contribute specificity, architecture, material properties and ligand-accessible regulatory nodes (Gilat et al., 2026; Peran and Mittag, 2020; Mohanty et al., 2022; Hess and Joseph, 2025; Wang and Marrink, 2026).

At the isolated-IDR1 level, S22 phosphorylation produced an apparent decoupling between dilute-chain dimensions and phase-separation propensity. For many IDRs, changes in single-chain dimensions provide useful information about the balance of intramolecular and intermolecular interactions. Here, however, phospho-IDR1 sampled slightly more expanded single-chain conformations than WT-IDR1, yet showed stronger phase separation in direct-coexistence simulations. Thus, phosphorylation-enhanced IDR1 condensation is not explained by compaction of the dilute- chain ensemble. Instead, dense-phase contact maps show that the added phosphate converts the S22/D26-containing region into a stronger anionic contact node that engages Arg- and Lys-rich patches on other chains. The salt dependence of these phosphorylation-enhanced contacts further supports an electrostatic origin for this rewiring. S22 phosphorylation therefore increases IDR1 phase-separation tendency by changing the specificity and distribution of interchain contacts in the dense phase, rather than by globally increasing single-chain collapse.

The IDR1 ligand response extends current models of small-molecule regulation of phase separation. Small molecules and other non-scaffold ligands are increasingly recognized as condensate regulators that can dissolve, induce, stabilize, relocalize or alter the material properties of biomolecular condensates (Mitrea et al., 2022; Patel et al., 2022; Usman et al., 2025; Kilgore and Young, 2022). Theory and simulation have further shown that ligand effects on scaffold phase behavior depend not only on ligand partitioning, but also on ligand valence, whether the ligand binds sticker-like or spacer-like sites, and the relative strengths of ligand–scaffold versus scaffold–scaffold interactions (Ruff et al., 2021). In this framework, ligands can destabilize condensates by competing with native scaffold–scaffold stickers, stabilize condensates by creating alternative crosslinks, or alter condensate organization even when their effects on phase stability are modest. Our IDR1 simulations provide a molecular example of this principle for two HDAC6-targeting ligands that were not originally designed as condensate modulators.

Both Nexturastat A and HPOB preferentially associate with positively charged IDR1 regions, positioning the ligands to compete with native acidic-to-basic contacts that stabilize the condensate. However, the two ligands redistribute this charged network differently. Nexturastat A uses a more focused polar-contact mode to engage Arg/Lys-rich patches and strongly depletes phospho- S22/D26-to-basic-patch contacts, thereby weakening phospho-IDR1 condensation. HPOB also contacts cationic patches, but its more distributed polar-contact pattern allows alternative charged contacts to be preserved or formed. This compensatory behavior explains why HPOB has a weaker disruptive effect on phospho-IDR1 and can even strengthen the weaker WT-IDR1 condensate. Thus, the IDR1 simulations support a competition-and-compensation model in which ligand effects depend on how each compound redistributes the specific charged contacts that maintain the dense phase.

This state-dependent ligand response is particularly relevant in light of previous work showing that small molecules can act as biphasic or context-dependent modulators of protein liquid– liquid phase separation (Babinchak et al., 2020). In such systems, the same chemical species can promote condensation under one regime and suppress it under another, depending on concentration, charge balance, interaction valence and the pre-existing scaffold network. Our simulations were not designed to map complete ligand-concentration phase diagrams, but they reveal a related principle: the effect of a ligand depends on the interaction network into which it is introduced. HPOB strengthens WT-IDR1, where the native condensate lacks the strong phospho- S22-centered anionic node, but weakens phospho-IDR1, where the ligand partially competes with a more specific phosphorylation-reinforced acidic-to-basic network. Nexturastat A, by contrast, acts more consistently as a focused competitor for cationic IDR1 patches. These findings argue that condensate pharmacology should not be interpreted solely in terms of ligand enrichment in the dense phase. Instead, ligand-induced changes in *C*_sat_, contact specificity, network connectivity and contact lifetime need to be considered together.

The IDR1-level mechanism, however, is not sufficient to explain full-length HDAC6. In full-length phospho-HDAC6 assemblies, the dominant contact feature was not an IDR1-centered network but FD3–FD3 association. FD3 corresponds to the C-terminal ZnF-UBP folded domain, indicating that a folded-domain surface contributes directly to the multivalent architecture of the assembled state. Moreover, S22 phosphorylation shifted the full-length assembly toward stronger FD3-centered organization, while several IDR1-associated contacts were reduced. These results show that the effect of phosphorylation is propagated through the multidomain architecture of HDAC6 rather than remaining confined to the modified IDR. More generally, they emphasize that mechanisms inferred from isolated disordered segments cannot be assumed to transfer directly to full-length multidomain proteins, where folded surfaces, domain connectivity and intervening disordered linkers reshape the accessible interaction network.

This conclusion is consistent with recent and emerging studies of multidomain condensates. Systematic mapping of IDR–IDR interactions suggests that many condensate-forming IDRs can coassemble promiscuously, implying that discrete condensate identity often requires additional structural determinants (Gilat et al., 2026). Folded domains can provide such determinants by contributing stereospecific surfaces, domain–motif interactions, oligomerization interfaces, nucleic-acid-binding modules or structured scaffolding elements (Li et al., 2012; Peran and Mittag, 2020; Mohanty et al., 2022; Hess and Joseph, 2025). Recent molecular simulations further suggest that folded domains can impose structural heterogeneity and attenuated dynamics in multidomain condensates, acting as architectural keystones rather than passive cargo (Wang and Marrink, 2026). Our FD3-centered phospho-HDAC6 assemblies provide a specific example of this principle: the folded ZnF-UBP domain functions as a major organizer of the full-length contact network, even though the experimentally identified regulatory phosphorylation site lies within IDR1.

Pocket-binding ligands revealed an additional domain-level regulatory mechanism. In full-length phospho-HDAC6, both Nexturastat A and HPOB engaged the catalytic-domain pockets and increased contacts between the combined pocket regions and FD3. These pocket–FD3 contacts were accompanied by reduced FD3–FD3 association, indicating competition between two folded-domain contact modes. The effect was strongest for Nexturastat A: although Nexturastat A and HPOB produced similar average pocket–FD3 contact numbers, Nexturastat A showed higher pocket occupancy, longer pocket residence time, longer pocket–FD3 residence time and a competition index shifted closest to zero. Therefore, the stronger remodeling by Nexturastat A is not explained simply by forming more pocket–FD3 contacts on average. Instead, Nexturastat A appears to generate more persistent pocket-mediated competition against the original FD3–FD3 association mode.

These results broaden the concept of condensate-modifying ligands in two ways. First, they show that a ligand can regulate an IDR condensate by competing with or compensating specific charged contacts, rather than by nonspecifically partitioning into the dense phase. Second, they show that a ligand originally developed to bind a folded catalytic pocket can remodel a full-length condensate assembly by redirecting folded-domain contacts. Thus, pocket-binding ligands can act through both an IDR-facing mechanism and a folded-domain-facing mechanism. In HDAC6, Nexturastat A more strongly disrupts the phospho-S22/D26-to-Arg/Lys contact network in IDR1 and more persistently redirects the FD3-centered full-length network toward pocket–FD3 contacts. HPOB produces weaker disruption at both levels, consistent with its more compensatory IDR1 contact pattern and less persistent pocket-mediated FD3 redirection. This dual mechanism provides a molecular explanation for why Nexturastat A more strongly suppresses phospho-IDR1 and phospho-HDAC6 condensation than HPOB in the motivating experiments (Lu et al., 2024).

More generally, our findings suggest that small molecules do not need to dissolve condensates by globally suppressing all interchain contacts. Instead, they can alter condensate organization by reweighting competing multivalent contact modes. In IDR1, this reweighting occurs among charged residue patches. In full-length phospho-HDAC6, it occurs between FD3–FD3 and pocket–FD3 domain contacts. This view is conceptually close to ligand-linkage models of phase separation, in which ligands modulate condensate stability and organization by changing the effective connectivity of scaffold interaction networks (Ruff et al., 2021). The HDAC6 system adds an important multidomain- protein extension to this framework: the relevant ligand-sensitive sites need not be located within the IDR that initially promotes condensation, but can reside in folded domains that redirect the architecture of the full-length assembly.

This model generates several testable predictions. Mutations that neutralize the phospho- S22/D26-centered acidic region or weaken the Arg/Lys-rich IDR1 patches should reduce the phosphorylation-dependent enhancement of IDR1 condensation and diminish IDR1-level ligand sensitivity. Conversely, mutations on the FD3 surface that weaken FD3–FD3 association should alter full-length HDAC6 assembly organization and reduce the extent to which catalytic-pocket ligands can redirect the FD3-centered network. Pocket mutations that preserve folded-domain stability but impair ligand engagement should reduce pocket–FD3 redirection without necessarily eliminating IDR1-mediated effects. Ligand analogs that separate pocket residence from IDR charge-patch association would be particularly informative: analogs retaining pocket binding but losing IDR charge-patch contacts should preferentially affect the full-length domain network, whereas analogs retaining IDR charge-patch engagement but showing weaker pocket residence should preferentially affect isolated IDR1 condensates. These predictions could be tested by combining mutational analysis, condensate imaging, turbidity or partitioning assays, fluorescence recovery measurements, crosslinking mass spectrometry and higher-resolution simulations.

Several considerations define the scope of the present work. First, the full-length simulations used finite 15-chain systems and were designed to resolve contact-network organization within associated multichain assemblies. They should therefore be interpreted as mechanistic analyses of domain-level reorganization, not as quantitative phase diagrams or full thermodynamic measurements of phase-separation propensity. Second, the coarse-grained framework enabled microsecond-scale sampling of large multidomain assemblies, but necessarily reduced atomistic detail. In particular, the simulations do not resolve explicit solvent structure, detailed side-chain packing, metal-coordination chemistry in the catalytic pockets, atomistic hydrogen-bond directionality or precise ligand-binding free energies. The FD2–ligand validation simulations therefore establish qualitative pocket-recognition behavior rather than quantitative affinity. Third, the cellular environment contains additional regulatory factors, including HDAC6 binding partners, ubiquitin chains, microtubules, molecular chaperones, additional post-translational modifications and macromolecular crowding. These factors may further tune the balance between IDR-mediated and folded-domain-mediated interactions in cells.

## Conclusions

Overall, our simulations support a two-layer mechanism for phospho-HDAC6 condensate regulation. At the IDR level, S22 phosphorylation enhances IDR1 phase separation by creating dense-phase electrostatic contacts between the phospho-S22/D26-containing region and cationic IDR1 patches. At the full-length protein level, however, the assembled network is dominated by FD3-centered folded-domain contacts rather than by IDR1 alone. This separation between the phosphorylation-responsive IDR layer and the FD3-centered folded-domain layer explains why isolated-IDR behavior cannot be directly extrapolated to the full multidomain protein.

The ligand response reveals a chemical route for controlling this multidomain assembly. Nexturastat A and HPOB do not simply suppress all condensate-forming contacts; instead, they redistribute competing interaction modes. In IDR1 condensates, they remodel charged residue-level contacts, whereas in full-length phospho-HDAC6 they redirect the folded-domain network from FD3–FD3 association toward pocket–FD3 contacts. The stronger and longer-lived remodeling by Nexturastat A shows that conventional catalytic-pocket ligands can act as contact-network modulators of condensate architecture. These findings suggest that ligand-accessible folded domains can be exploited as chemical control points for multidomain-protein condensates, providing a framework for designing pocket-targeting molecules that regulate phase-separated assemblies by reweighting residue- and domain-level interactions.

## Materials and Methods

### Protein model construction

The starting structural model of human HDAC6 was obtained from the AlphaFold Protein Structure Database entry AF-Q9UBN7 (Jumper et al., 2021; Varadi et al., 2022). Before topology generation, the per-residue pLDDT profile was used to define folded and disordered blocks in the full-length protein (Figure S2A). Contiguous regions with overall pLDDT values above 90 were assigned as folded domains, using this conservative high-confidence cutoff to avoid treating poorly resolved regions as structured blocks. Short lower-pLDDT segments embedded within high-confidence domains were retained within the corresponding folded domain and stabilized with Gō-type restraints, rather than being separated into additional disordered segments. Low-confidence regions connecting high-pLDDT blocks were assigned as IDR-like regions.

This pLDDT-guided assignment defined six blocks for model construction, visualization and downstream contact analysis: IDR1, residues 1–85; FD1, residues 86–438; IDR-L, residues 439–480; FD2, residues 481–835; IDR2, residues 836–1111; and FD3, residues 1112–1215. The isolated-IDR1 simulations were designed to match the experimentally defined IDR1 construct and therefore used residues 1–66, whereas the full-length HDAC6 model retained the pLDDT-defined IDR1 block spanning residues 1–85.

The full-length HDAC6 coarse-grained topology was generated using Martinize2 (Kroon et al., 2025). Martini3-IDP was used as the base representation for disordered protein regions (Wang et al., 2025). GōMartini 3 restraints were assigned to the folded regions FD1, residues 86–438; FD2, residues 481–835; and the folded core of FD3, residues 1112–1208 (Souza et al., 2021, 2025). The remaining IDR-like regions, residues 1–85, 439–480, 836–1111 and the C-terminal segment 1209–1215, were left flexible under the Martini3-IDP representation. Thus, the model preserved the structural integrity of high-confidence folded domains while allowing the disordered regions and terminal low-confidence segments to sample flexible conformations.

Because S22 phosphorylation is central to the HDAC6 condensate phenotype, we constructed a local phosphoserine parameter set for the S22 site before production simulations. A previously reported Martini 3 phosphoserine approximation was used as the starting point (Linhartova et al., 2024). The S22 side-chain bead was assigned as Q5 with charge −1*e*, while the remaining protein bead assignment was kept unchanged. Local bonded terms around Gln21, phospho-Ser22 and Pro23 were then calibrated against mapped atomistic reference trajectories. In the fitted local model, bead 44 is the Gln21 backbone bead (P2), bead 45 is the Gln21 side-chain bead (P5), bead 46 is the Ser22 backbone bead (P2), bead 47 is the phosphorylated Ser22 side-chain bead (Q5, charge−1*e*), bead 48 is the Pro23 backbone bead (SP2a), and bead 49 is the Pro23 side-chain bead (SC3). The fitted bonded terms were selected to capture both the internal phosphoserine geometry and the coupling between S22 and its neighboring residues. These terms included the 46–47 S22 backbone–side-chain bond; the 44–46–47 and 47–46–48 angles linking S22 to Gln21 and Pro23; the 45–44–46–47 and 47–46–48–49 dihedrals spanning the adjacent side chains; and the 46–44–48–47 improper dihedral describing the local three-residue geometry.

Agreement between the mapped atomistic-reference distribution and the calibrated coarse-grained distribution was quantified for each fitted term using the Hellinger distance,

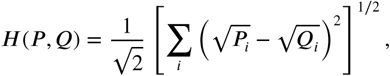

where *P_i_* and *Q_i_* are normalized histogram probabilities for the atomistic-reference and coarse-grained validation distributions, respectively. The six Hellinger distances were low to moderate across the fitted terms, with the largest value observed for the 47–46–48–49 dihedral (Figure S2B). Distribution-level comparisons for all six fitted terms are shown in Figure S2C–H.

To reduce the known sensitivity of coarse-grained IDR phase behavior to the balance of protein– water interactions, we applied a uniform IDR–water environmental-bias correction adapted from the GōMartini 3 strategy (Souza et al., 2025). In the original implementation, an additional environmental-bias term between virtual Gō sites in strand residues and water beads was introduced with an energy parameter of −0.5 kJ mol^−1^. We used the same bias strength for the IDR–water correction and applied it uniformly to simulations involving the IDR models.

### Small-molecule coarse-grained model construction

Nexturastat A and HPOB were represented as Martini 3 coarse-grained ligands following established Martini protein–ligand and small-molecule modeling strategies (Souza et al., 2020; Alessandri et al., 2022). Atomistic structures of Nexturastat A and HPOB were extracted from PDB entries 5G0I and 5EF7, respectively (Hai and Christianson, 2016; Miyake et al., 2016). These structures provided the initial small-molecule geometries for coarse-grained model construction. Both ligands were treated as neutral molecules.

Initial atom-to-bead mappings and Martini 3 bead types were generated using Automartini_M3 (Szczuka et al., 2026). The bonded geometry of each ligand was then refined against mapped atomistic ligand trajectories using PyCGTOOL (Graham et al., 2017), yielding GROMACS-compatible topology files for production simulations. The final Nexturastat A model contained 12 neutral beads, comprising seven C-type beads, four N-type beads and one P-type bead. The final HPOB model contained 11 neutral beads, comprising six C-type beads, three N-type beads and two P-type beads. Final atom-to-bead mappings are shown in Figure 1B, and the same ligand color convention is used throughout all figures involving ligand representations.

### Molecular dynamics simulations

Atomistic systems were prepared with CHARMM-GUI (Jo et al., 2008; Lee et al., 2016) and simulated using the CHARMM36m force field (Huang et al., 2017) with TIP3P water (Jorgensen et al., 1983).

Periodic boundary conditions were applied in all three dimensions. Energy minimization was performed using the steepest-descent algorithm for up to 5000 steps, with an energy tolerance of 1000 kJ mol^−1^ nm^−1^. Systems were equilibrated for 125 ps with a 1 fs timestep and then simulated for 500 ns with a 2 fs timestep. Bonds involving hydrogen atoms were constrained using LINCS (Hess et al., 1997). Van der Waals interactions used a 1.2 nm cutoff, and long-range electrostatics were treated using particle-mesh Ewald electrostatics (Darden et al., 1993; Essmann et al., 1995) with a 1.2 nm real-space cutoff. Temperature was maintained at 295 K using the velocity-rescaling thermostat (Bussi et al., 2007) with a 1 ps coupling time constant. Production simulations were performed under isotropic NPT conditions at 1 bar using the C-rescale barostat (Bernetti and Bussi, 2020), with a 5 ps pressure-coupling time constant and a compressibility of 4.5 × 10^−5^ bar^−1^.

Atomistic simulations were used to generate reference distributions for local phosphoserine bonded-term calibration and ligand bonded-geometry refinement. Atomistic single-chain simulations of the residues 1–66 IDR1 construct were performed in 15 × 15 × 15 nm^3^ boxes at 50 mM NaCl for three independent 500 ns replicates. The maximum intramolecular atomic distance was approximately 11.1 nm, indicating that periodic-image effects were avoided. Isolated Nexturastat A and HPOB simulations were performed in three independent 500 ns replicates per ligand.

Coarse-grained simulations used the Martini 3-based protein and ligand models described above, with Martini3-IDP representations for disordered regions and GōMartini 3 restraints for folded domains (Souza et al., 2021; Wang et al., 2025; Souza et al., 2025). Periodic boundary conditions were applied in all three dimensions, and all coarse-grained simulations were performed at 295 K. Energy minimization used the steepest-descent algorithm for up to 200,000 steps. Equilibration used the Berendsen barostat (Berendsen et al., 1984), and production simulations used a 20 fs timestep with the Parrinello–Rahman barostat (Parrinello and Rahman, 1981). The Verlet cutoff scheme was used with neighbor-list updates every 20 steps and a Verlet buffer tolerance of 0.005 kJ mol^−1^ ps^−1^ (Verlet, 1967; Páll and Hess, 2013). Electrostatic interactions were treated with the reaction-field method using a relative dielectric constant of 15 and a 1.1 nm cutoff (Tironi et al., 1995). Van der Waals interactions used a 1.1 nm cutoff with the potential-shift-Verlet scheme. Temperature was maintained using the velocity-rescaling thermostat with a 1 ps coupling time constant (Bussi et al., 2007). Pressure was maintained at 1 bar with a 12 ps pressure-coupling time constant. Slab simulations used semi-isotropic pressure coupling with fixed lateral box dimensions, whereas all other coarse-grained systems used isotropic pressure coupling. No constraints were applied in the coarse-grained simulations.

Full-length HDAC6 simulations were initialized in 30 × 30 × 30 nm^3^ boxes containing 15 HDAC6 chains at 50 mM NaCl, with additional counterions added where necessary to ensure charge neutrality. Ligand-treated systems additionally contained 50 molecules of either Nexturastat A or HPOB, corresponding to an initial ligand concentration of 3075.1 *µ*M. Each full-length condition was simulated for three independent 10 *µ*s replicates.

Coarse-grained single-chain simulations of IDR1 used the residues 1–66 construct and were performed in 20 × 20 × 20 nm^3^ boxes at 50 mM NaCl for three independent 20 *µ*s replicates. Direct-coexistence slab simulations used the same residues 1–66 IDR1 construct and were initialized in 20 × 20 × 50 nm^3^ boxes containing 150 IDR1 chains (Dignon et al., 2018; Mammen Regy et al., 2021; Benayad et al., 2021). The 20 nm lateral dimensions were chosen to reduce periodic-image effects, because the longest single-chain dimension was approximately 17.5 nm.

Initial slab configurations were generated using gmx insert-molecules, followed by slow compression of the chains toward the box center along the *z* direction using an inverted flat-bottomed potential under NVT conditions. The compressed systems were further equilibrated for 1 *µ*s while maintaining the inverted flat-bottomed potential, yielding an initial slab thickness of approximately 15 nm. Ligand-free slab simulations were performed at 0, 50 and 150 mM NaCl, with additional counterions added where necessary to ensure charge neutrality. Ligand-treated slab simulations were performed at 50 mM NaCl and contained 50 molecules of either Nexturastat A or HPOB, corresponding to an initial ligand concentration of 4151.4 *µ*M. IDR1 direct-coexistence slab simulations were run for 15 *µ*s and analyzed over the 11–15 *µ*s window after late-time density-profile convergence was assessed. Each IDR1 slab condition was simulated as one production replicate.

Dedicated FD2–ligand validation simulations contained one FD2 domain and one ligand molecule at 50 mM NaCl. Ten independent 10 *µ*s replicates were performed for each ligand. These simulations were used as qualitative pocket-recognition controls and were not interpreted as binding free-energy calculations.

All molecular dynamics simulations were performed with GROMACS 2021.7 (Abraham et al., 2015). Molecular configurations and trajectories were inspected using Visual Molecular Dynamics (VMD) (Humphrey et al., 1996). A complete overview of the atomistic and coarse-grained simulation systems, including box dimensions, compositions, salt concentrations, replicate numbers and production lengths, is provided in Table S2.

### Data analysis and statistics

Trajectory analyses were performed using GROMACS tools and custom Python scripts. For IDR1 slab simulations, protein density profiles were calculated from mass density along the slab-normal *s* axis using gmx density with 100 slabs, centering, relative coordinates and symmetric averaging. Production density profiles were calculated from the 11–15 *µ*s analysis window for ligand-free and ligand-treated slabs. Apparent dilute-phase concentrations were obtained from the same density profiles using the custom Python workflow used to generate the slab-density plots.

For each density profile, the workflow calculated *z*_max_ = max(|*z*|) from the density-profile coordinate array. The dense-phase region was defined as bins satisfying |*z*| ≤ 0.20*s*_max_, and the dilute-phase region was defined as bins satisfying |*z*| ≥ 0.80*s*_max_. If either mask was empty, the workflow used the highest-density 10% of bins as the dense-phase fallback or the lowest-density 10% of bins as the dilute-phase fallback. Dense- and dilute-phase mass densities were calculated as arithmetic means of the density bins in the corresponding masks, with missing values ignored. The apparent dilute-phase concentration reported in the figures was the dilute-region mean density, expressed in mg ml^−1^. For these profiles, kg m^−3^ is numerically equivalent to mg ml^−1^. The plotted uncertainty was the sample standard deviation of the density bins in the dilute-phase mask, using *n* − 1 normalization when more than one dilute bin was available and zero otherwise.

For residues 1–66 IDR1 single-chain simulations, the radius of gyration was calculated using gmx gyrate. The end-to-end distance was calculated as the distance between the centers of mass of residues 1 and 66. Pair-distance distribution functions were calculated from all pairwise distances among protein beads. For each replicate, pair-distance histograms were accumulated over frames and normalized to probability density. Replicate-normalized curves were then averaged across the three independent simulations. Detailed histogram settings are provided in the caption of Figure S5C.

Dense-phase IDR1 contact maps were calculated from the largest IDR1-rich cluster in each slab trajectory over the 11–15 *µ*s production window. The dense-phase cluster was identified using the DBSCAN clustering algorithm (Ester et al., 1996), with periodic boundary conditions applied to the distance calculations. Two IDR1 chains were considered connected when any interchain bead–bead distance was below 0.60 nm, and the largest cluster satisfying the minimum cluster-size criterion of three chains was retained for contact analysis. Only chains in this largest dense-phase cluster were used for residue-level contact maps. Interchain residue–residue contacts within the cluster were accumulated over analyzed frames using a 0.60 nm bead–bead cutoff and converted to contact-probability maps.

Ligand–IDR1 contacts in ligand-treated slab simulations were calculated over the 11–15 *µ*s analysis window using periodic minimum-image distances and a 0.60 nm bead–bead cutoff. A residue–ligand contact was counted when at least one bead from a given protein residue was within 0.60 nm of a ligand bead. Contacts with positively charged residues were calculated by restricting the protein residue set to Arg and Lys. Contacts with negatively charged residues were calculated by restricting the protein residue set to Asp, Glu and phospho-Ser residues.

Dense-phase ligand molecules were defined as ligand molecules whose centroids were located within the *z*-extent of the largest IDR1-rich cluster. Ligand-bead-resolved contact maps were calculated between dense-phase ligand molecules and residues belonging to the largest IDR1-rich cluster. Ligand-centered radial number-density profiles were calculated from periodic minimum-image distances between dense-phase ligand centroids and charged-residue centroids. Radial shells were normalized by shell volume and by the number of dense-phase ligand centroids.

For full-length HDAC6 simulations, time-resolved interchain contact numbers were calculated using gmx mindist between different HDAC6 chains with a 0.60 nm bead–bead cutoff and periodic boundary conditions. Domain-level contact maps used the pLDDT-defined domain blocks: IDR1, residues 1–85; FD1, residues 86–438; IDR-L, residues 439–480; FD2, residues 481–835; IDR2, residues 836–1111; and FD3, residues 1112–1215. Interchain bead–bead contacts within 0.60 nm were assigned to domain pairs. Domain-level maps were calculated over the final 3 *µ*s analysis window, normalized by the number of analyzed frames, interchain chain pairs and domain bead counts, and then averaged across independent replicates.

Pocket analyses in full-length HDAC6 simulations used two catalytic-pocket definitions mapped onto the full-length HDAC6 sequence. Pocket residues were derived from inhibitor-bound HDAC6 catalytic-domain structures, including PDB entry 5G0I, by selecting residues within 1.0 nm of the co-crystallized ligand, mapping these residues onto the full-length HDAC6 sequence by sequence alignment, and retaining mapped positions only when residue identity matched the full-length model (Hai and Christianson, 2016; Miyake et al., 2016; Osko et al., 2020). Pocket1 comprised residues 99, 101, 102, 104, 106, 107, 111, 172–175, 212–217, 224, 226, 251, 253–258, 277, 283–286, 342–346, 351–354, and 382–386. Pocket2 comprised residues 494, 497–502, 506, 568, 569, 607– 612, 618–621, 647, 649–654, 679–681, 740–742, 747–750, and 778–782. For isolated FD2-domain validation simulations, the domain-local pocket residues were 15, 18–23, 27, 88–91, 128–133, 139–142, 168, 170–175, 199–203, 261–263, 268–271, and 299–303.

Ligand pocket occupancy was defined by a minimum ligand–pocket bead distance below 0.60 nm for either pocket1 or pocket2. In full-length simulations, mean pocket occupancy was averaged over pocket1 and pocket2, all 15 protein chains and all analyzed frames in each replicate. Binary pocket-occupancy traces were converted into residence events by grouping consecutive occupied frames. A one-frame interruption between two occupied intervals was treated as part of the same event. Residence time was measured from the first occupied frame to the last occupied frame, including one frame interval, such that a single occupied frame contributed one frame spacing. Mean residence times were obtained by pooling events over pocket1 and pocket2 and over all chains within each replicate before calculating replicate-level means. FD2–ligand validation contact frequencies were calculated as the fraction of frames in which the ligand contacted the FD2 pocket. For the 15-chain full-length HDAC6 simulations, pocket–FD3 contact numbers were calculated over the 7–10 *µ*s window. For each frame, interchain bead–bead contacts within 0.60 nm were counted between pocket1 or pocket2 residues on one chain and FD3 residues on another chain. FD3–FD3 contacts used the same cutoff and were restricted to interchain FD3 bead pairs. The competition index was calculated frame by frame as

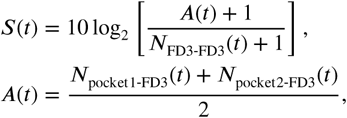

where larger values indicate a shift from FD3–FD3 contacts toward pocket–FD3 contacts. The additive pseudocount of 1 avoided division by zero, and the factor of 10 was used only to scale the plotted values and does not change the ordering of systems.

For the two-chain phospho-HDAC6 pairwise analysis shown in Figure S11, the same pocket–FD3, FD3–FD3 and competition-index definitions were applied to the 6–10 *µ*s window of the FD2-centered two-chain trajectories. This two-chain analysis was used to examine pairwise domain-interface reorganization and was not interpreted as a direct phase-separation assay.

FD3–pocket residence was calculated from binary engagement traces between each focal pocket- centered reference and partner-chain FD3. In ligand-free simulations, the reference object was the pocket itself. In ligand-treated simulations, all frames were analyzed as a single drug-system state. For each frame, the reference object was the pocket itself when no ligand bead was within 0.60 nm of that pocket. When the pocket was occupied, the reference object was defined as the union of the pocket and the seeded ligand cluster. Seed molecules were ligand molecules with at least one bead within 0.60 nm of the pocket, and the seeded cluster was expanded to include ligand molecules connected to a seed by ligand–ligand bead contacts within 0.60 nm. Residence outputs were not further stratified by ligand-occupied and ligand-unoccupied states.

FD3 engagement was defined by the minimum distance between a partner-chain FD3 and the pocket-centered reference. A frame was classified as engaged when this distance was below 1.00 nm. Residence events were defined as continuous engaged intervals for a given focal reference and partner-chain FD3 pair. Event duration was calculated as the time from the first engaged frame to the last engaged frame plus the 2 ns frame spacing. No additional short-gap merging was applied.

When independent simulations were available, each independent simulation replicate was treated as the statistical unit. Scalar quantities from full-length simulations are reported as mean ± s.e.m., where s.e.m. denotes the standard error of the mean, across three independent replicates. FD2–ligand validation contact frequencies are reported as mean ± s.e.m. across ten independent replicates. IDR1 single-chain radius-of-gyration and end-to-end-distance summaries are reported as mean ± s.e.m. across three independent replicates. IDR1 slab density profiles, dense-phase contact maps, ligand-contact decompositions and ligand-centered radial-density pro-files were calculated from single production slab trajectories and are therefore shown without replicate-level s.e.m. For time-resolved full-length contact traces, representative traces are shown in the main text and replicate-level traces are provided in the Supporting Information where available.

## Supplementary files

Supplementary File 1. Additional simulation details, model construction, parameterization, analysis procedures, supplementary figures, and supplementary tables (PDF).

## Funding

X.C. acknowledges support from the National Natural Science Foundation of China (Grant No. 12474201), the Guangdong Provincial Project (Grant No. 2023QN10X037), and the Guangdong S&T Program (Grant No. 2025A0505000027).

## Acknowledgments

The authors acknowledge the Green e Materials (GeM) Laboratory and the HPC+AI Intelligence Computing Center at the Hong Kong University of Science and Technology (Guangzhou) for providing computational resources.

## Competing interests

The authors declare no competing financial interest.

## Data Availability

The source data supporting the main-text and Supplementary Information figures and analyses are available in the GitHub repository (https://github.com/cygooo/HDAC6-phosphorylation-pocket-ligand-condensate-networks/ tree/main/source_data). The repository includes a file-level index in source_data/README.md that maps each deposited file or file pattern to the corresponding figure, analysis output, or validation dataset.

## Code Availability

Custom scripts used for data processing, quantity calculation and figure generation are available in the GitHub repository (https://github.com/cygooo/HDAC6-phosphorylation-pocket-ligand-condensate-networks/ tree/main/code_availability). The repository includes a script-level index in code_availability/README.md that maps each script to the corresponding calculation or plotting task.

---

**Figure S1:**
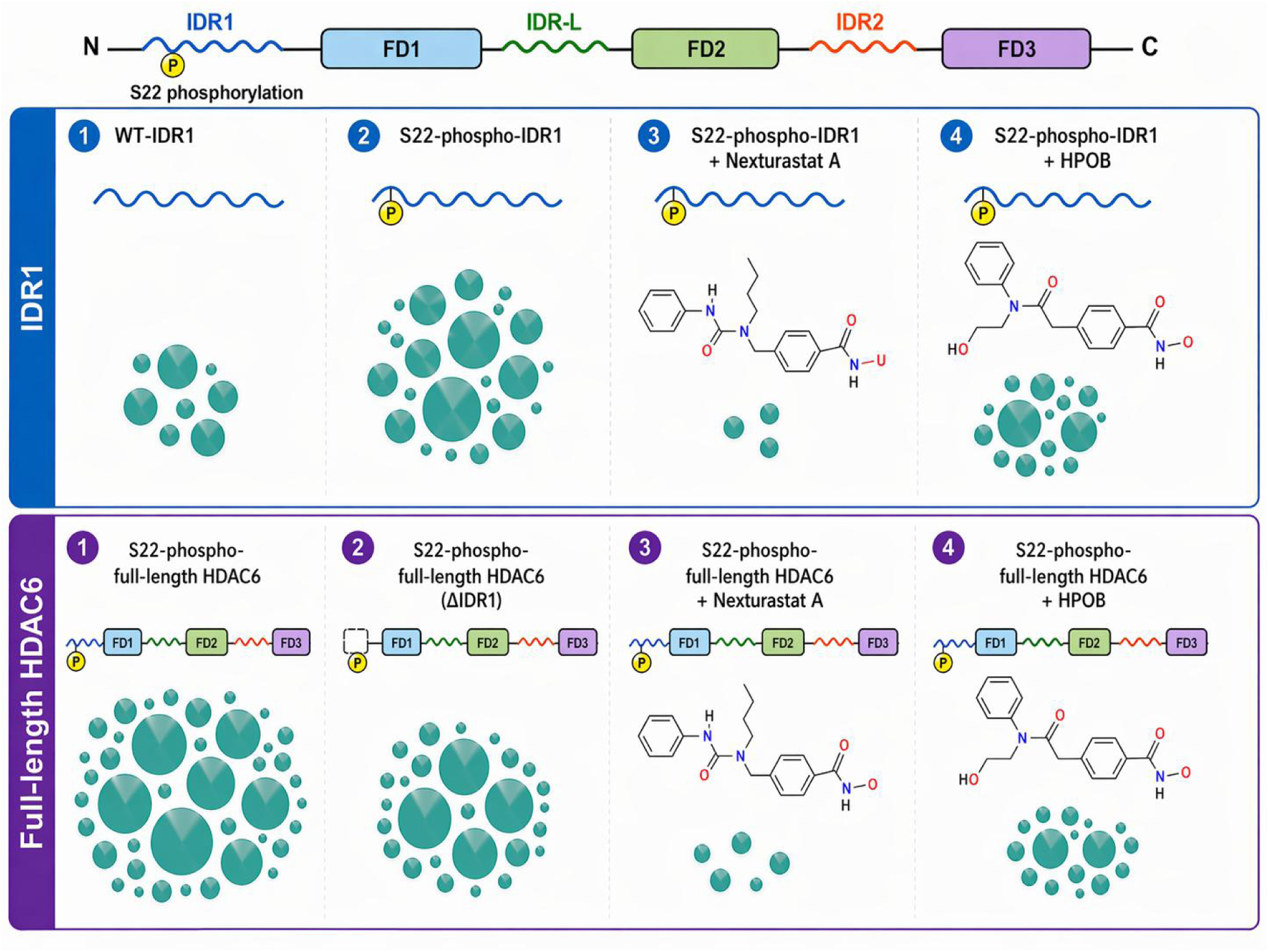
Schematic summary of experimentally reported HDAC6 phase-separation phenotypes motivating the present simulation study. The top schematic shows the modular organization of HDAC6, including the N-terminal disordered region IDR1, folded domains FD1 and FD2, the linker disordered region IDR-L, IDR2, and the C-terminal folded domain FD3. The upper row summarizes IDR1-level phenotypes: S22 phosphorylation enhances the phase-separation tendency of IDR1 relative to WT-IDR1, Nexturastat A strongly suppresses S22-phospho-IDR1 condensation, and HPOB has a weaker suppressive effect. The lower row summarizes full-length HDAC6 phenotypes: S22-phosphorylated full-length HDAC6 forms condensates, removal of IDR1 reduces but does not abolish condensation of the remaining HDAC6 construct, Nexturastat A strongly suppresses full-length phospho-HDAC6 condensation, and HPOB has a weaker suppressive effect. This schematic summarizes previously reported experimental phenotypes and is not a quantitative representation of condensate size or number.

**Figure S2:**
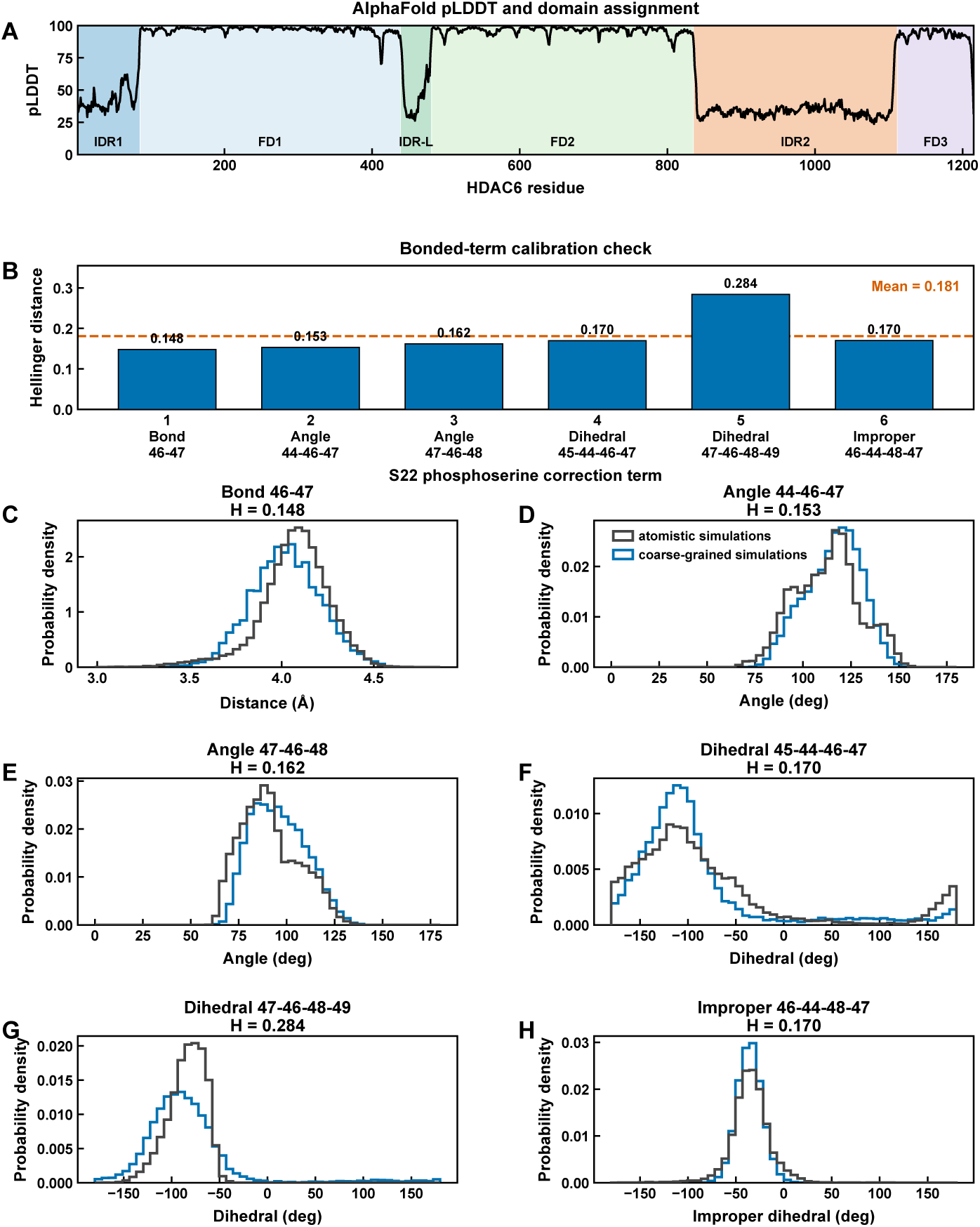
Protein model construction and S22 phosphoserine parameter validation. **(A)** Per-residue pLDDT profile from the AlphaFold HDAC6 model used for coarse-grained model construction. Colored blocks mark the domain definitions used for model construction and analysis: IDR1, FD1, IDR-L, FD2, IDR2 and FD3. **(B)** Hellinger distances comparing mapped atomistic-reference distributions with calibrated coarse-grained distributions for six local bonded terms around phospho-S22, including one bond, two angles, two dihedrals and one improper dihedral. The dashed line indicates the mean Hellinger distance across the six terms. **(C–H)** Distribution-level comparisons between atomistic-reference simulations and calibrated coarse-grained simulations for the six local phospho-S22 bonded terms shown in **B**. Black and blue curves indicate atomistic-reference and coarse-grained distributions, respectively.

**Figure S3:**
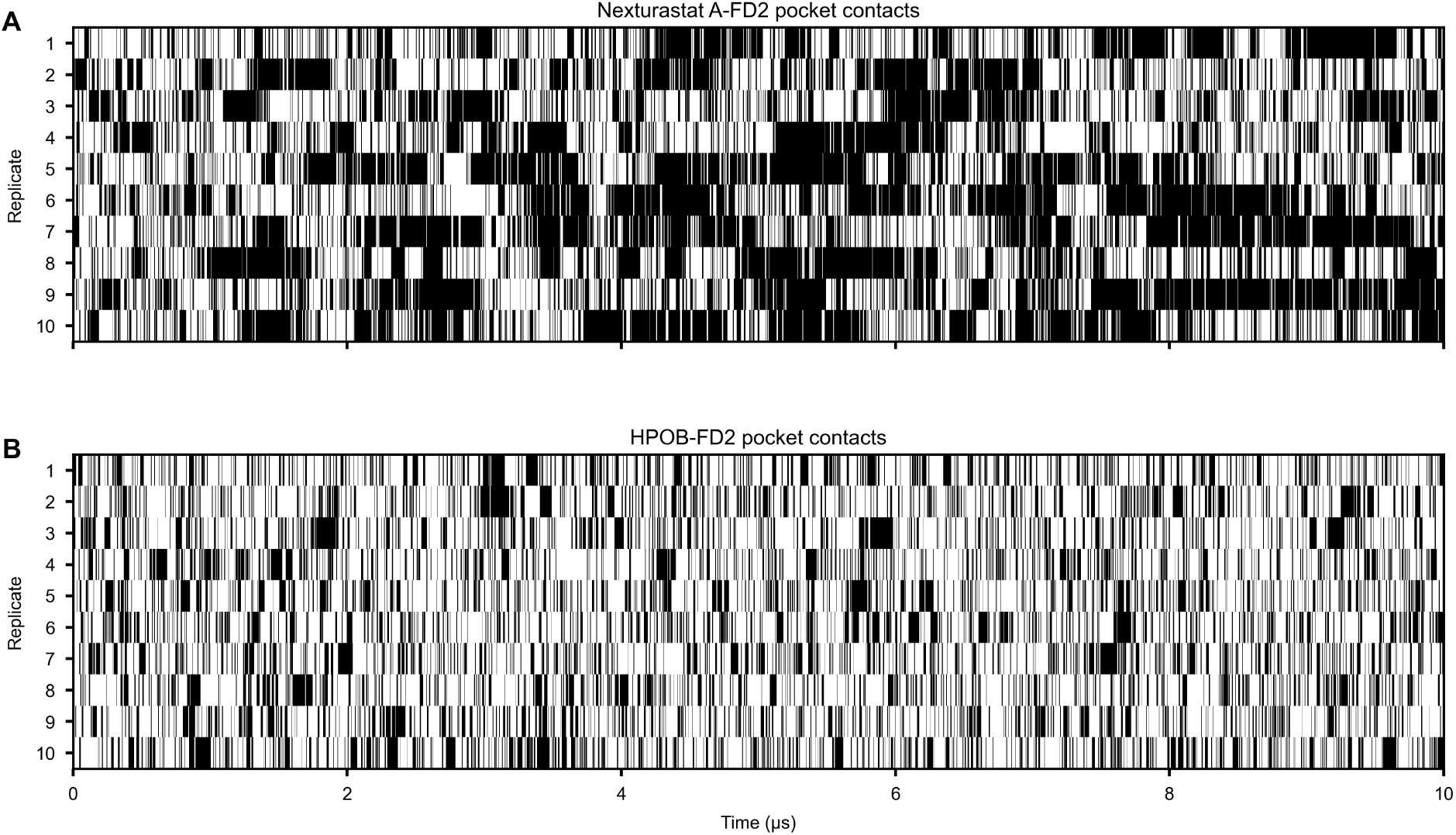
Time-resolved ligand–pocket contact barcode plots from dedicated one-ligand validation simulations. **(A)** Nexturastat A–FD2 pocket contacts across ten independent 10 *µ*s simulations. **(B)** HPOB–FD2 pocket contacts across ten independent 10 *µ*s simulations. In each panel, each row represents one replicate simulation and the horizontal axis shows simulation time. Black marks indicate frames in which the ligand contacts the FD2 pocket, whereas white regions indicate no pocket contact, using the same contact definition as in the contact-fraction analysis in Fig. 1C.

**Figure S4:**
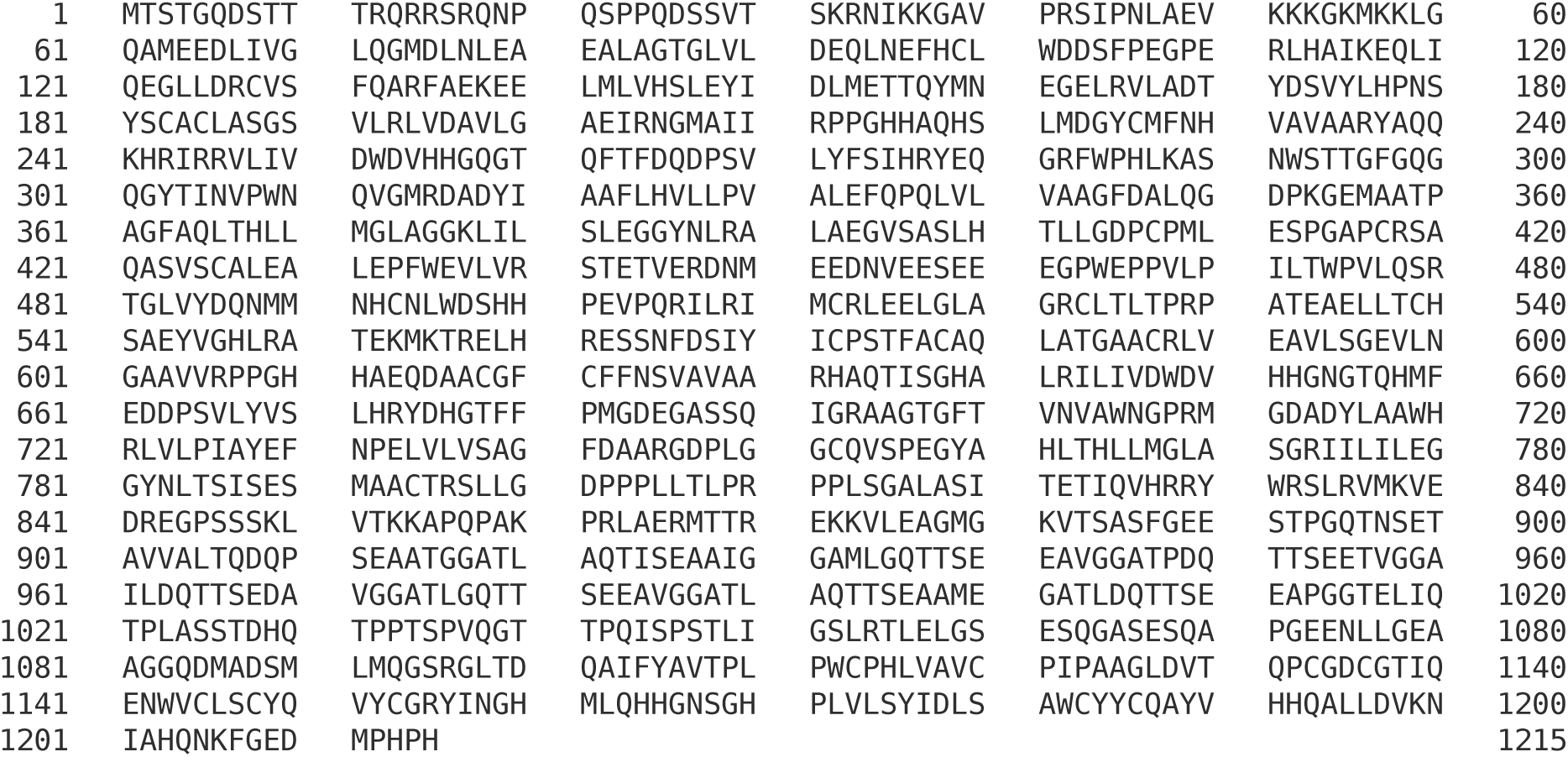
Human full-length HDAC6 sequence used for model construction. The sequence used to construct the full-length HDAC6 coarse-grained model was obtained from the AlphaFold Protein Structure Database entry AF-Q9UBN7 and is shown with residue numbering. Residues are grouped in blocks of ten amino acids for readability.

**Figure S5:**
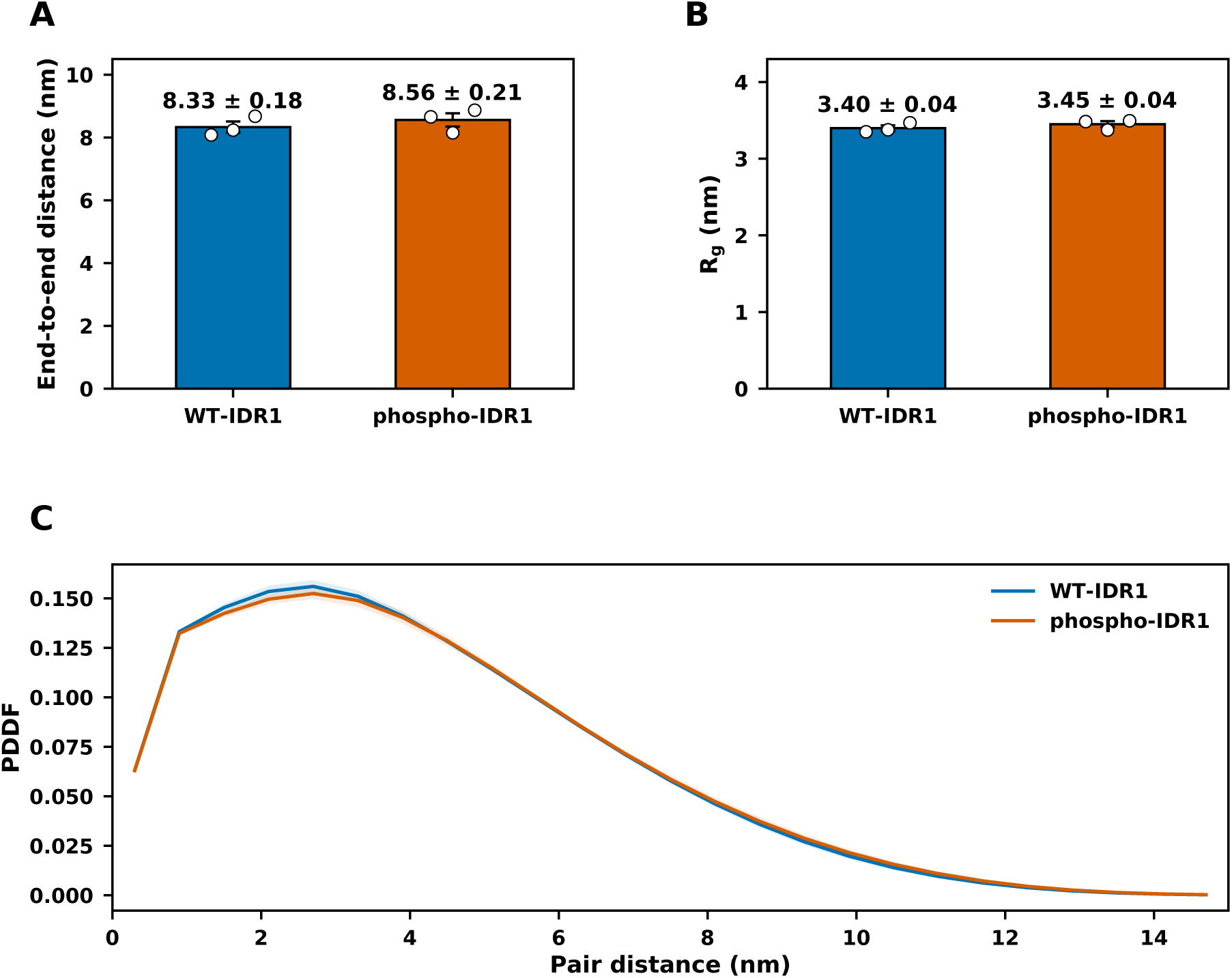
Single-chain conformational metrics for WT-IDR1 and phospho-IDR1 at 50 mM NaCl. **(A,B)** End-to-end distance and radius of gyration (*R_g_*) for single-chain simulations of the residues 1–66 WT-IDR1 and phospho-IDR1 constructs. Bars show mean values, error bars indicate s.e.m., and open circles indicate means from individual simulations. **(C)** Intrachain pair-distance distribution functions (PDDFs) calculated from all pairwise protein-bead distances, normalized to probability density and averaged across three independent simulations.

**Figure S6:**
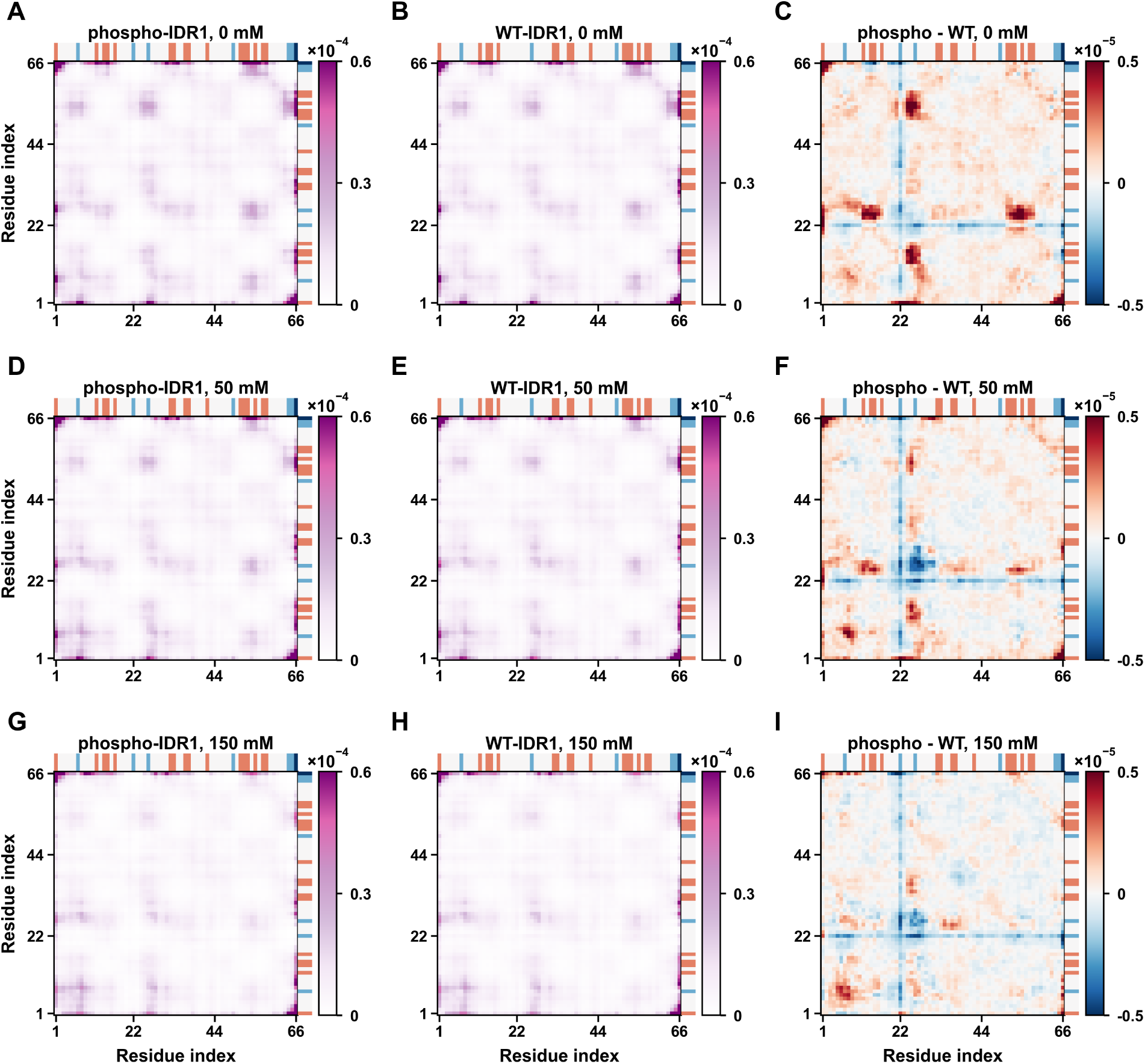
Salt-dependent dense-phase interchain contact maps of WT-IDR1 and phospho-IDR1. Dense-phase interchain residue–residue contact probability maps from WT-IDR1 and phospho-IDR1 slab simulations at 0, 50 and 150 mM NaCl, calculated over the 11–15 *µ*s analysis window. **(A,D,G)** Contact maps of phospho-IDR1 at 0, 50 and 150 mM NaCl, respectively. **(B,E,H)** Contact maps of WT-IDR1 at 0, 50 and 150 mM NaCl, respectively. **(C,F,I)** Difference maps calculated as phospho-IDR1 minus WT-IDR1 at the corresponding salt concentrations. Red regions indicate contacts enhanced by S22 phosphorylation, whereas blue regions indicate contacts reduced by phosphorylation. Charge strips above and to the right of each map indicate residue-level charge states using a discrete encoding: red, +1; white, 0; light blue, -1; dark blue, -2. Phosphorylation-enhanced contacts involving the S22-containing region and positively charged IDR1 clusters are strongest at low salt and become weaker at higher salt, supporting an electrostatic contribution to dense-phase contact rewiring.

**Figure S7:**
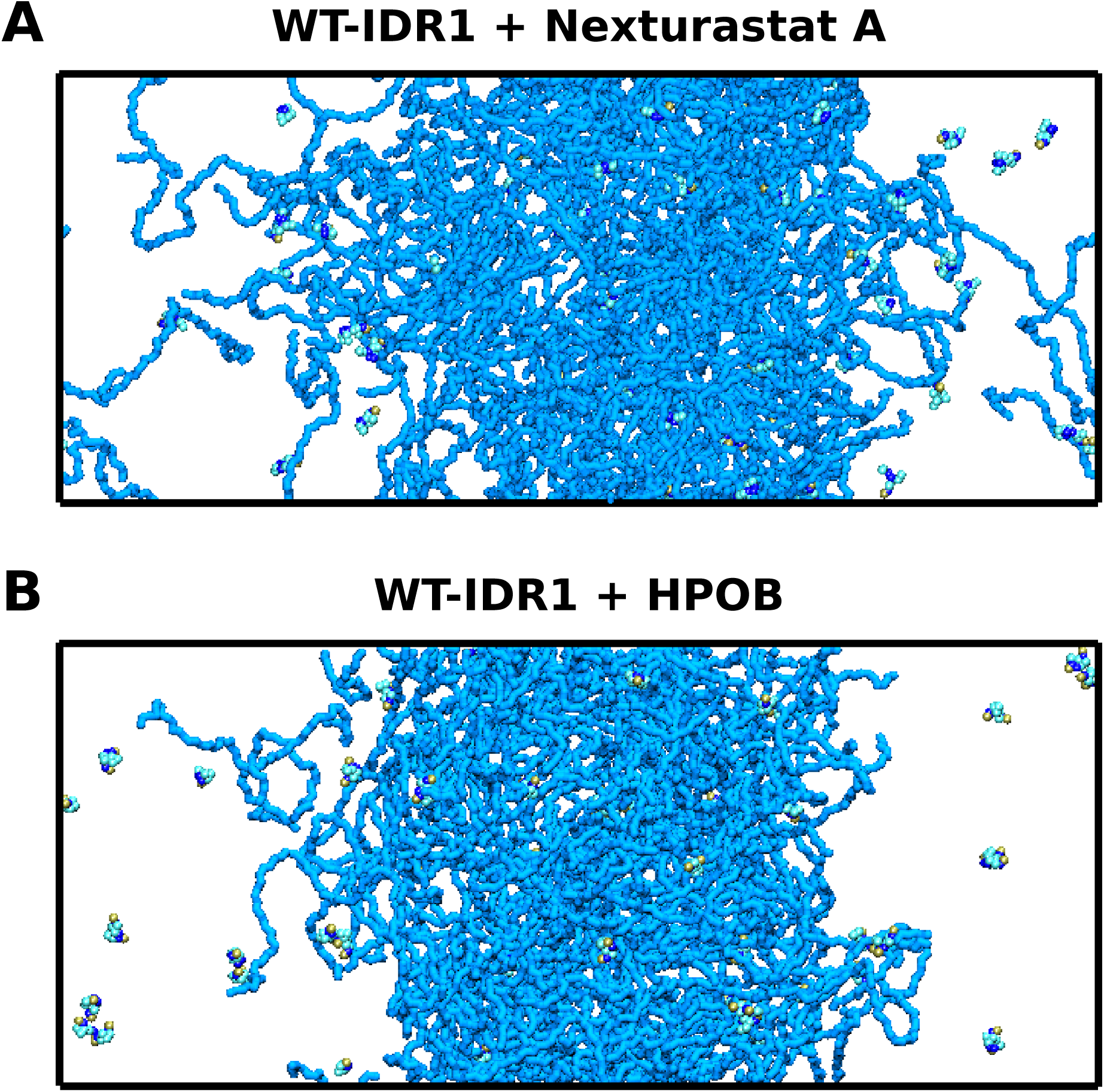
Representative ligand-treated WT-IDR1 slab snapshots. **(A,B)** Representative snapshots taken at 15 *µ*s from WT-IDR1 slab simulations treated with Nexturastat A and HPOB, respectively, at 50 mM NaCl. Snapshots were cropped consistently and displayed at matched scale.

**Figure S8:**
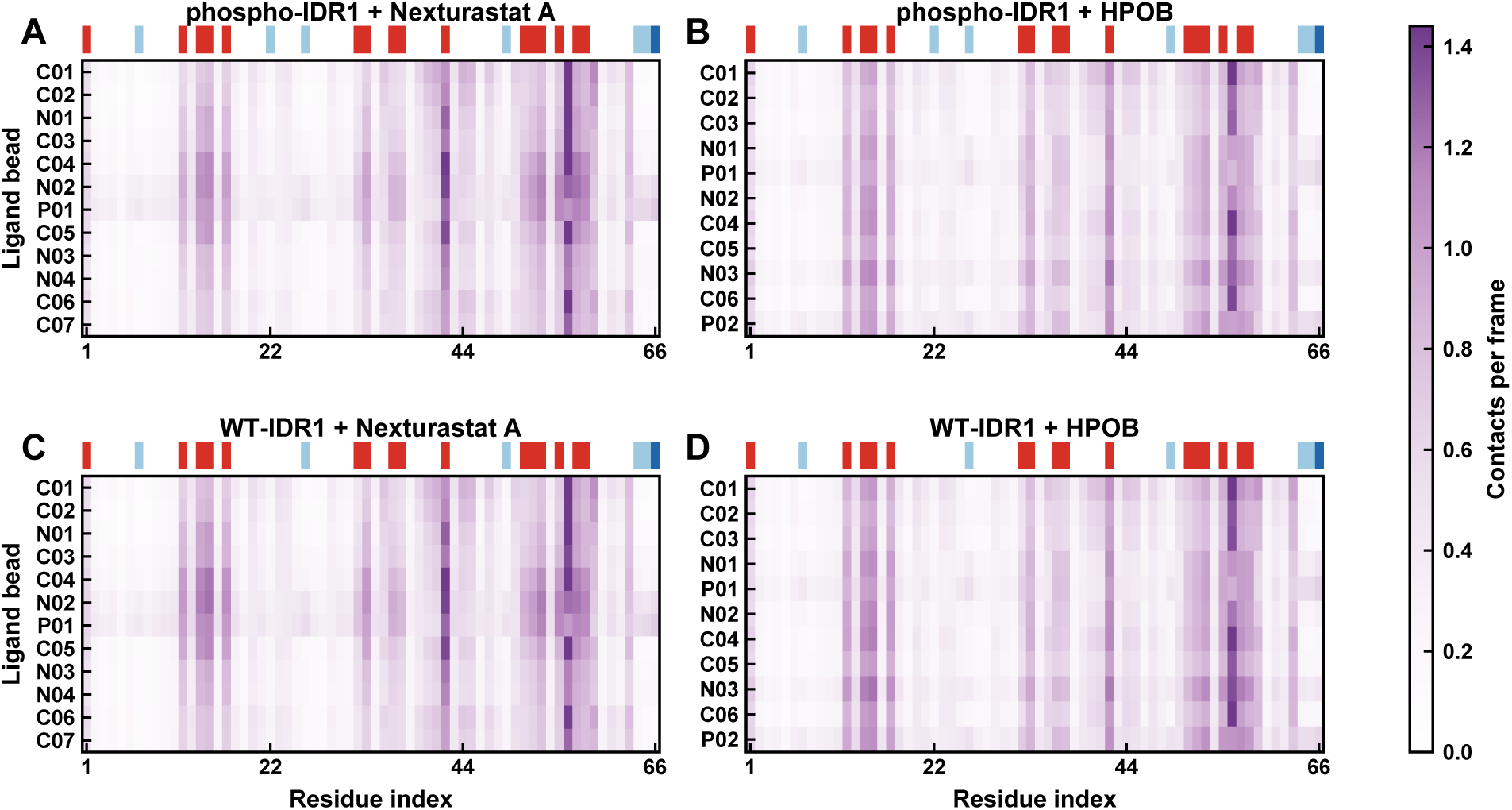
Ligand-bead-resolved contact maps between IDR1 and dense-phase ligand molecules. **(A,B)** Ligand-bead versus residue contact maps for phospho-IDR1 treated with Nexturastat A and HPOB, respectively. **(C,D)** Ligand-bead versus residue contact maps for WTIDR1 treated with Nexturastat A and HPOB, respectively. Contacts were calculated from the dense-phase regions of slab simulations at 50 mM NaCl over the 11–15 *µ*s analysis window. All panels use the same color scale. Charge strips above each map indicate residue-level charge states using a discrete encoding: red, +1; white, 0; light blue, -1; dark blue, -2. The maps indicate that ligand contacts are concentrated around positively charged IDR1 regions, with bead-level patterns that are consistent with a more localized polar-head contribution for Nexturastat A and a broader polar-contact contribution for HPOB. Strong bead-level signals involve donor- or acceptor-subtype polar beads, supporting a coarse-grained polar-contact interpretation rather than atomistically resolved directional hydrogen bonds.

**Figure S9:**
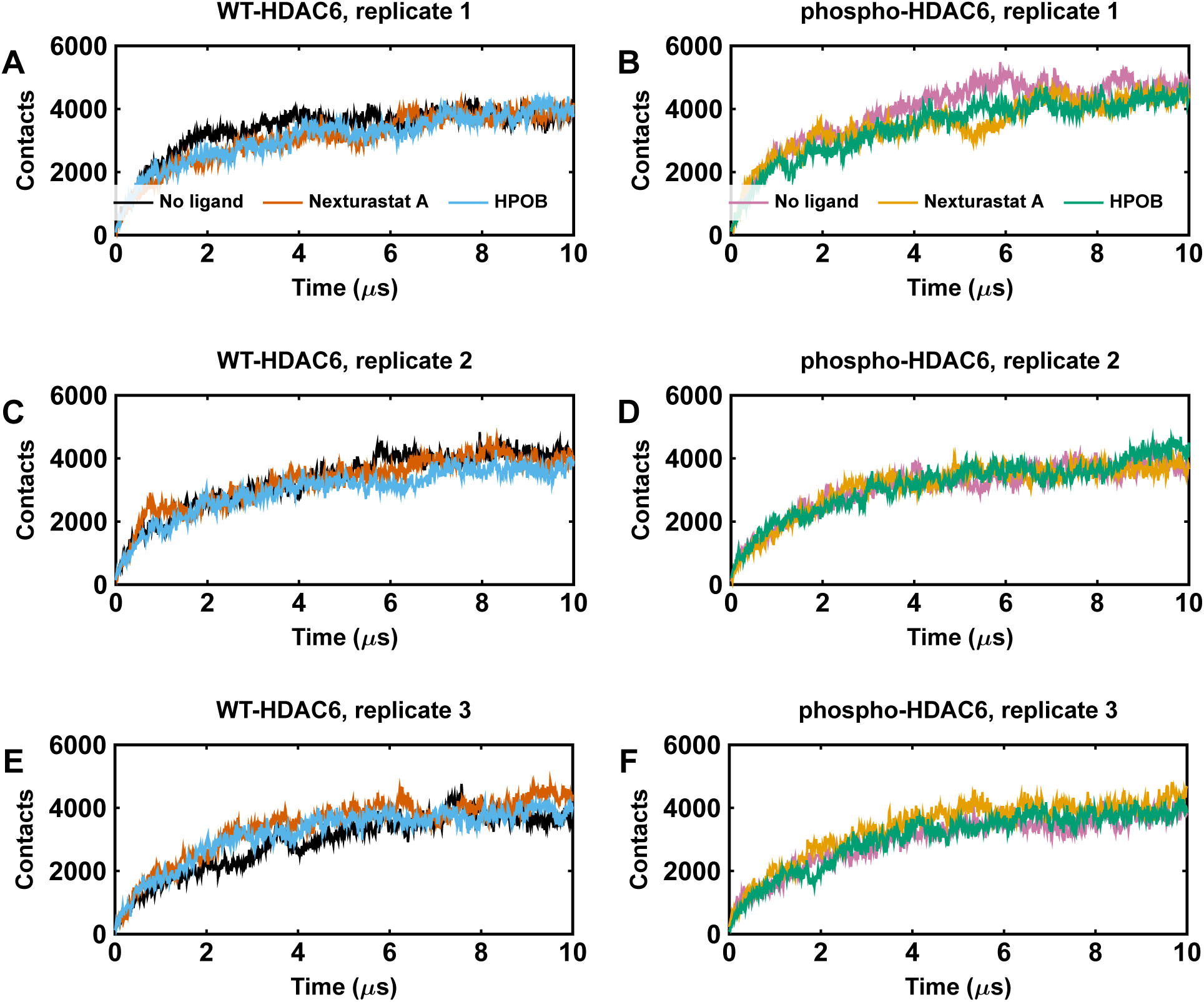
Replicate-level interchain contact-number traces from full-length HDAC6 simulations. Time-resolved interchain contact numbers are shown for three independent simulations of WT-HDAC6 and phospho-HDAC6 under no-ligand, Nexturastat A-treated and HPOB-treated conditions. **(A,C,E)** WT-HDAC6 replicates 1–3, respectively. **(B,D,F)** Phospho-HDAC6 replicates 1–3, respectively. Within each panel, traces compare the three ligand conditions over 10 *µ*s simulations. These replicate-level traces support the interchain contact comparison summarized in Fig. 4A.

**Figure S10:**
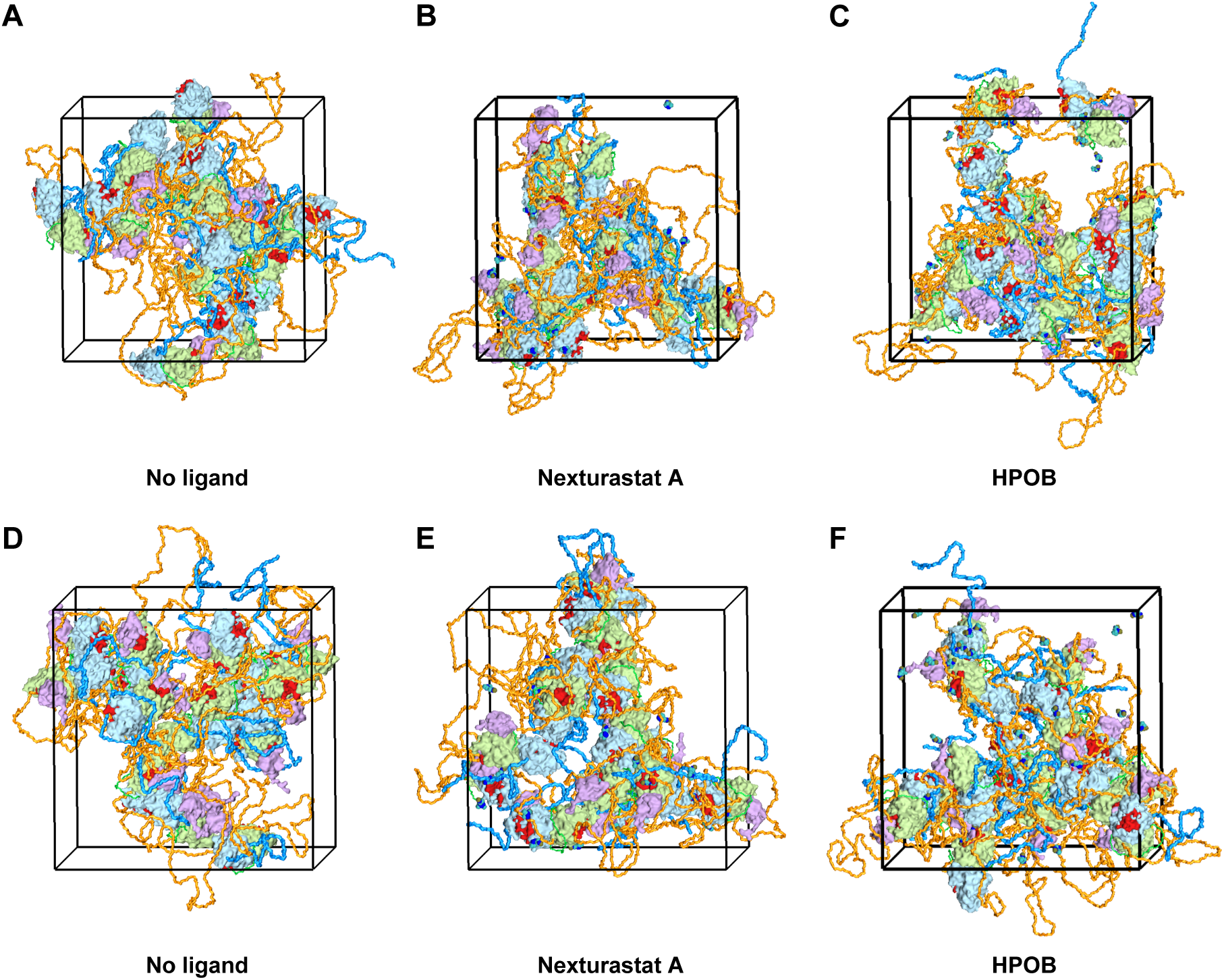
Representative full-length HDAC6 snapshots from 15-chain simulations. **(A–C)** Representative snapshots taken at 10 *µ*s from phospho-HDAC6 simulations under no-ligand, Nexturastat A-treated and HPOB-treated conditions, respectively. **(D–F)** Representative snapshots taken at 10 *µ*s from WT-HDAC6 simulations under no-ligand, Nexturastat A-treated and HPOB-treated conditions, respectively. FD1, FD2 and FD3 are shown as folded-domain surfaces, IDR segments are shown as chains, and catalytic-pocket regions are highlighted in red.

**Figure S11:**
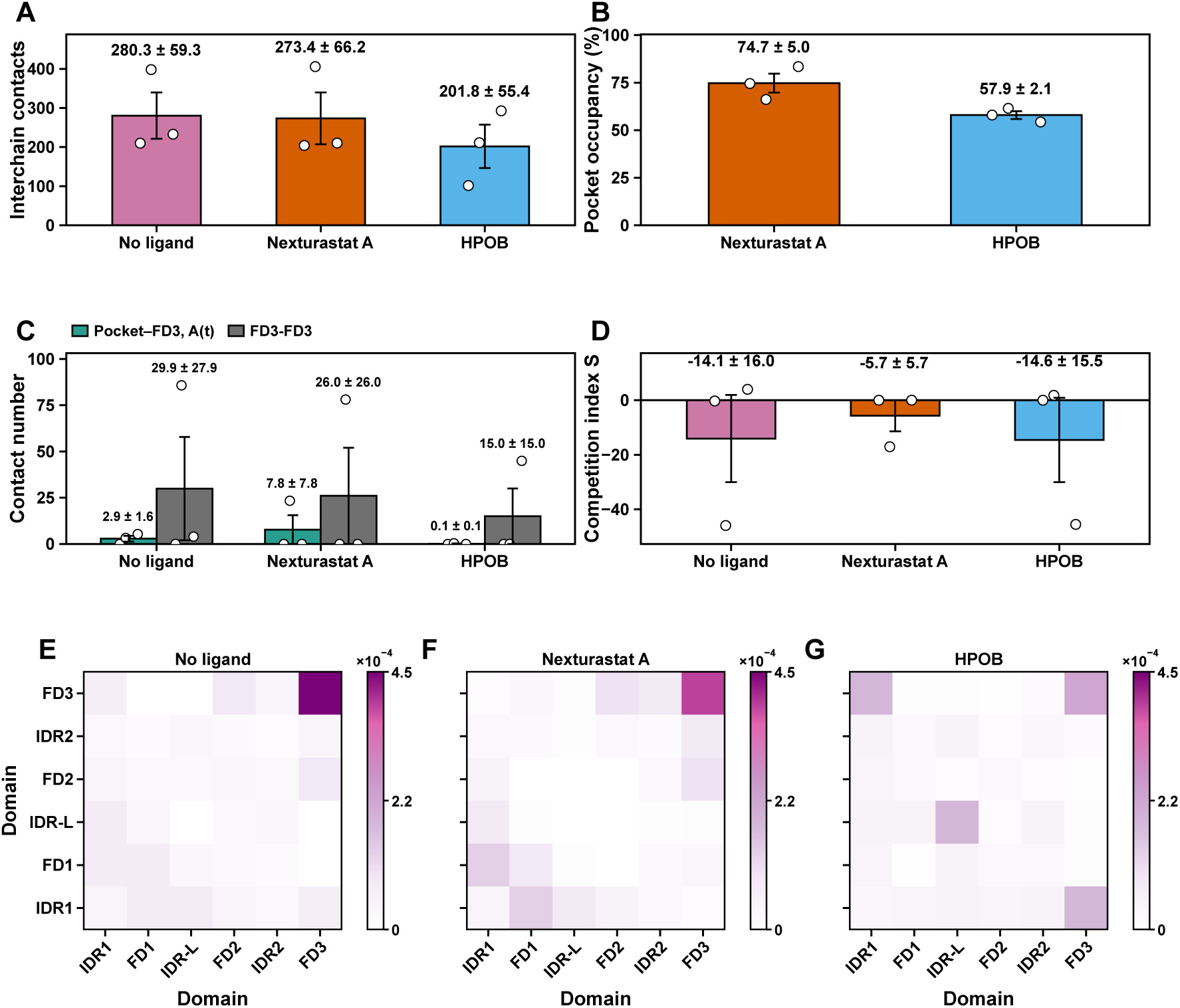
Pairwise domain-interface rewiring in two-chain phospho-HDAC6 simulations. Two-chain S22-phosphorylated full-length HDAC6 simulations were analyzed over the 6–10 *µ*s analysis window to examine pairwise domain-interface organization. These simulations were used to resolve pairwise contact rewiring rather than to quantify phase-separation propensity. **(A)** Mean total interchain contact number under no-ligand, Nexturastat A-treated and HPOB-treated conditions, showing that the two chains remained associated in all conditions. **(B)** Mean pocket occupancy for Nexturastat A and HPOB, showing ligand engagement with the catalytic-pocket regions. **(C)** Mean pocket–FD3 and FD3–FD3 contact numbers under no-ligand, Nexturastat A-treated and HPOB-treated conditions, used to calculate the competition index. **(D)** Competition index *S* under no-ligand, Nexturastat A-treated and HPOB-treated conditions, calculated from pocket–FD3 and FD3–FD3 contact numbers; more positive or less negative values indicate a relative shift from FD3–FD3 association toward pocket-centered FD3 recruitment. **(E–G)** Normalized symmetric domain-level interchain contact matrices for no-ligand, Nexturastat A-treated and HPOB-treated two-chain phospho-HDAC6 simulations, respectively, shown using the same color scale. Bars show mean values, error bars indicate s.e.m. across three independent simulations, and open circles indicate individual simulations. Together, these two-chain simulations support the pairwise domain-rewiring mechanism inferred from the 15-chain full-length phospho-HDAC6 simulations, with Nexturastat A showing stronger pocket-centered redirection than HPOB.

**Figure S12:**
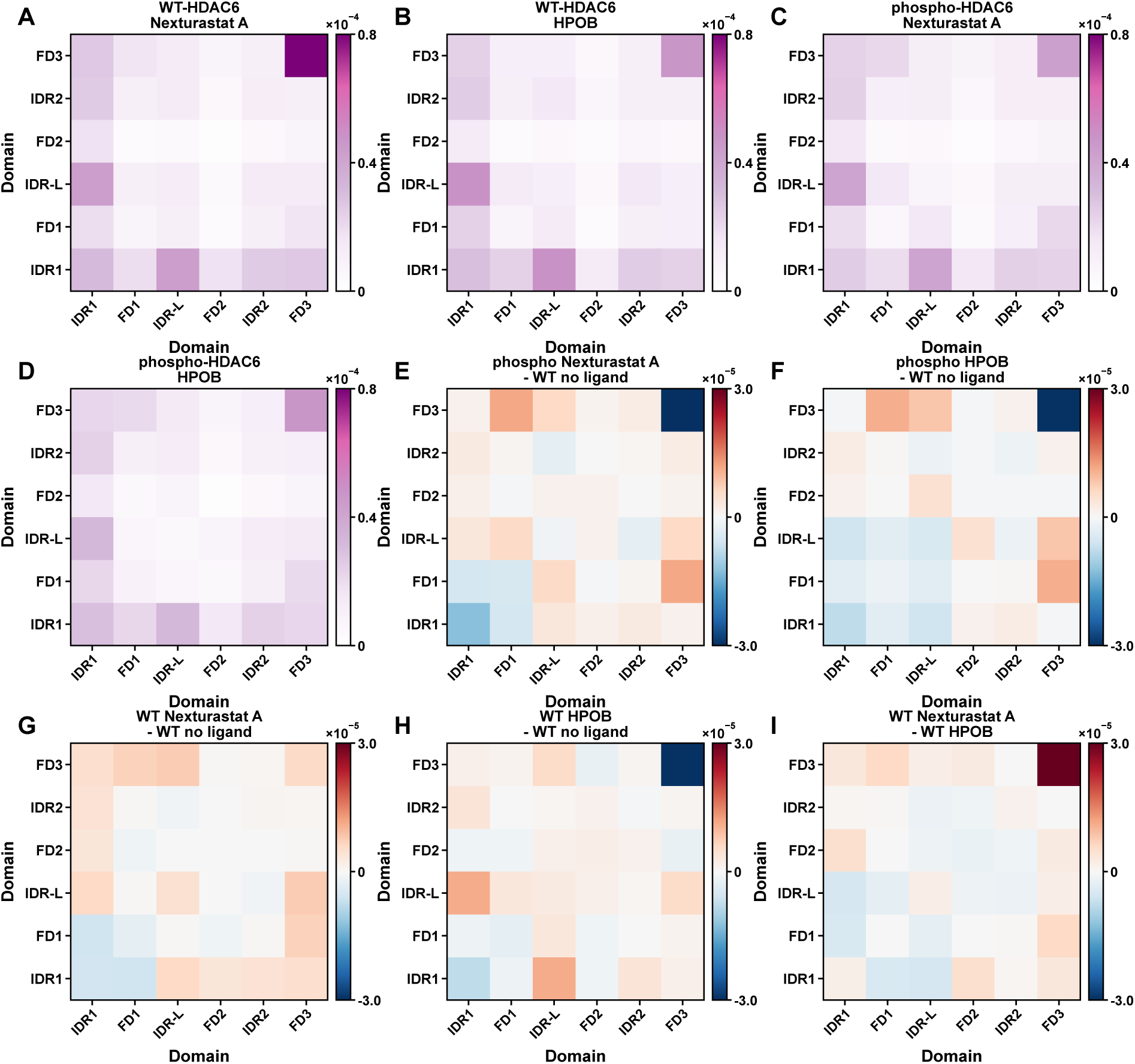
Supplementary full-length domain-level interchain contact maps and difference maps. Domain-level interchain contact probability maps and difference maps were calculated from the final 3 *µ*s of the 15-chain full-length HDAC6 simulations. **(A–D)** Domain-level interchain contact maps for WT-HDAC6 with Nexturastat A, WT-HDAC6 with HPOB, phospho-HDAC6 with Nexturastat A, and phospho-HDAC6 with HPOB, respectively. **(E,F)** Difference maps calculated as phospho-HDAC6 with Nexturastat A minus no-ligand WT-HDAC6 and phospho-HDAC6 with HPOB minus no-ligand WT-HDAC6, respectively. **(G,H)** Difference maps calculated as WT-HDAC6 with Nexturastat A minus no-ligand WT-HDAC6 and WT-HDAC6 with HPOB minus no-ligand WT-HDAC6, respectively. **(I)** Difference map calculated as WT-HDAC6 with Nexturastat A minus WT-HDAC6 with HPOB. Red indicates contacts increased in the first condition of each comparison, whereas blue indicates contacts reduced in the first condition. These panels complement the phospho-HDAC6-focused domain contact maps in Fig. 4B–G.

**Figure S13:**
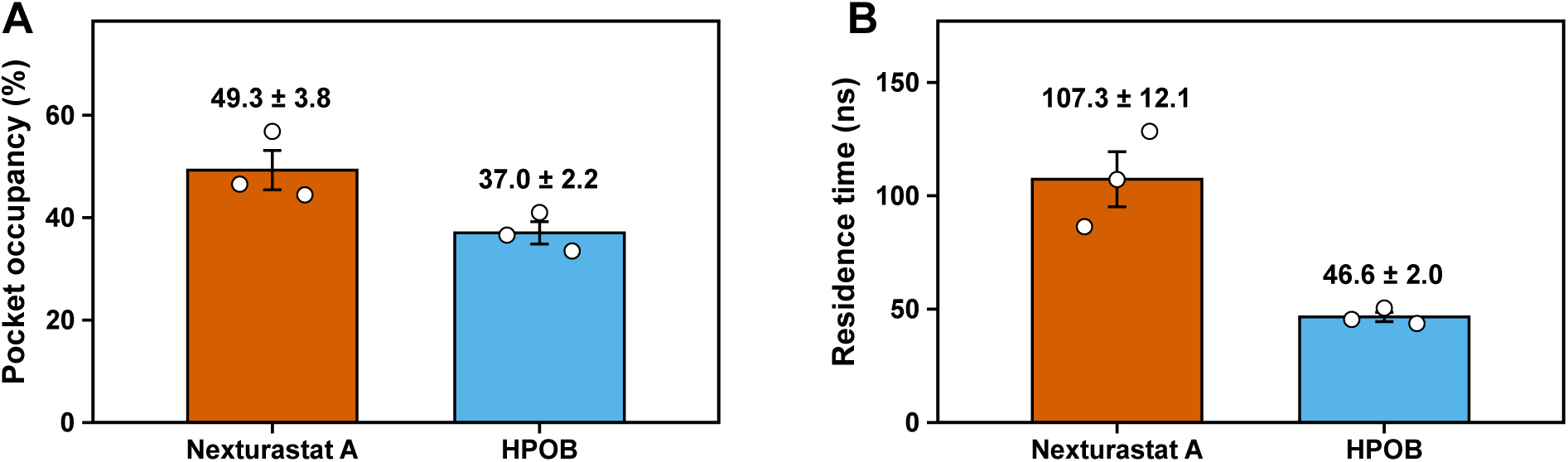
Pocket occupancy and residence time of Nexturastat A and HPOB in full-length WT-HDAC6 simulations. **(A)** Mean pocket occupancy for Nexturastat A and HPOB in full-length WT-HDAC6 simulations. **(B)** Mean pocket residence time for Nexturastat A and HPOB in the same simulations. Bars show mean values, error bars indicate s.e.m. across three independent simulations, and open circles indicate individual simulations.

**Figure S14:**
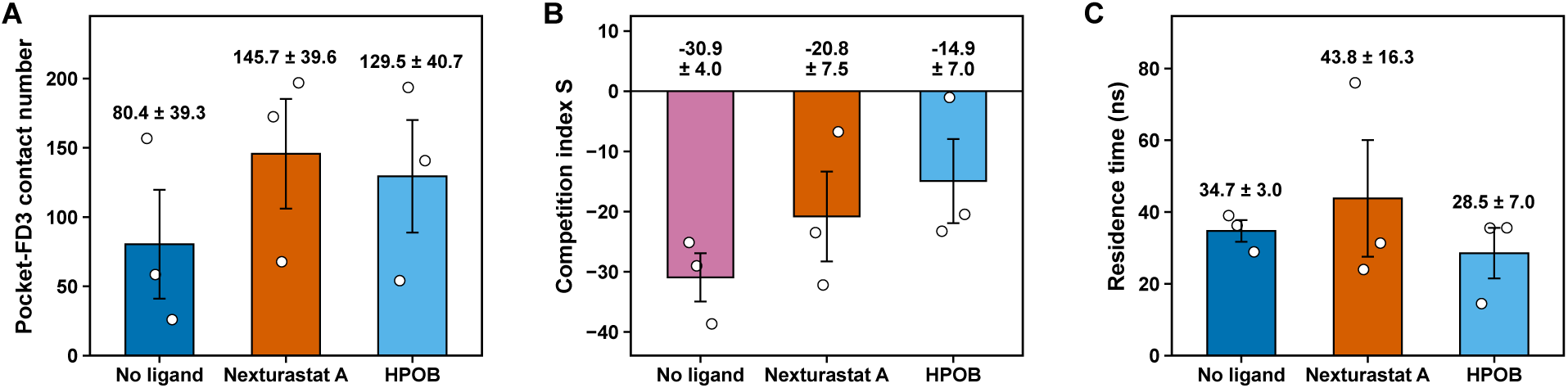
Pocket–FD3 contact metrics in full-length WT-HDAC6 simulations. **(A)** Mean pocket–FD3 contact number in WT-HDAC6 simulations under no-ligand, Nexturastat A-treated and HPOB-treated conditions. **(B)** Competition index *S*, calculated from pocket–FD3 and FD3–FD3 contact numbers for the same WT-HDAC6 simulations. More positive values indicate increased pocket–FD3 contacts relative to FD3–FD3 contacts. **(C)** Mean pocket–FD3 residence time in the corresponding WT-HDAC6 simulations. Bars show mean values, error bars indicate s.e.m. across three independent simulations, and open circles indicate individual simulations.

**Table S1:** Bead types used for the coarse-grained interpretation of ligand contacts with charged IDR1 residues. The assignments support interpretation of ligand–residue contacts as coarse-grained polar or electrostatic contacts. They do not define atomistically resolved directional hydrogen bonds.

| Molecule or residue | Bead label | Martini bead type | Contact role | Interpretation |
| --- | --- | --- | --- | --- |
| Arg side chain | SC2 | SQ3p | Positive residue contact partner | Cationic side-chain bead |
| Lys side chain | SC2 | SQ4p | Positive residue contact partner | Cationic side-chain bead |
| Asp side chain | SC1 | SQ5n | Negative residue contact partner | Anionic side-chain bead; the n subtype does not encode directional hydrogen bonding |
| Glu side chain | SC1 | Q5n | Negative residue contact partner | Anionic side-chain bead; the n subtype does not encode directional hydrogen bonding |
| phospho-Ser22 side chain | SC1 | Q5; $q = -1e$ | Negative residue contact partner | Charged phosphoserine bead |
| Nexturastat A | N02 | TN5a | Major Arg/Lys-contacting ligand bead | N5a polar bead with acceptor subtype at the polar headgroup |
| Nexturastat A | P01 | TP1d | Additional charged-residue-contacting ligand bead | P1d donor-subtype polar bead at the polar headgroup |
| HPOB | N03 | SN3a | Major Arg/Lys-contacting ligand bead | N3a polar bead with acceptor subtype at the hydroxamate/amide end |
| HPOB | P02 | TP1d | Hydroxamate/amide-end polar bead | P1d donor-subtype polar bead supporting coarse-grained polar contacts |
| HPOB | P01 | TP2d | Lateral hydroxyl-associated polar bead | P2d donor-subtype polar bead associated with the lateral hydroxyl group |

**Table S2:**
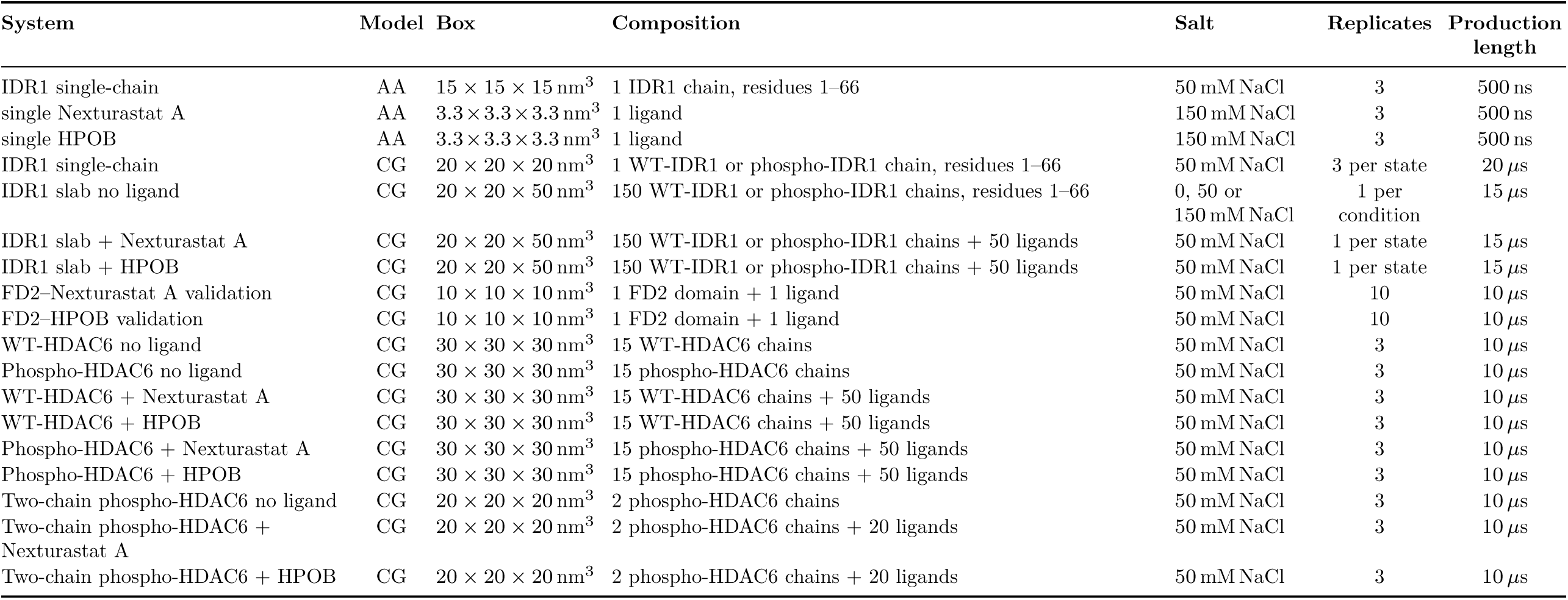
Summary of all-atom (AA) and coarse-grained (CG) simulations. The table lists the simulation systems used for atomistic reference calculations, ligand model refinement, coarse-grained validation and production analyses.

## Notes

### Competing Interest Statement

The authors have declared no competing interest.

